# Chromosome 9p21.3 Coordinates Cell Intrinsic and Extrinsic Tumor Suppression

**DOI:** 10.1101/2022.08.22.504793

**Authors:** Francisco M. Barriga, Kaloyan M. Tsanov, Yu-Jui Ho, Noor Sohail, Amy Zhang, Timour Baslan, Alexandra N. Wuest, Isabella Del Priore, Brigita Meškauskaitė, Geulah Livshits, Direna Alonso-Curbelo, Janelle Simon, Almudena Chaves-Perez, Dafna Bar-Sagi, Christine A. Iacobuzio-Donahue, Faiyaz Notta, Ronan Chaligne, Roshan Sharma, Dana Pe’er, Scott W. Lowe

## Abstract

Somatic chromosomal deletions are prevalent in cancer, yet their functional contributions remain ill-defined. Among the most prominent of these events are deletions of chromosome 9p21.3, which disable a cell intrinsic barrier to tumorigenesis by eliminating the *CDKN2A/B* tumor suppressor genes. However, half of 9p21.3 deletions encompass a cluster of 16 type I interferons (IFNs) whose co-deletions have not been functionally characterized. To dissect how 9p21.3 and other genomic deletions impact cancer, we developed MACHETE (Molecular Alteration of Chromosomes with Engineered Tandem Elements), a genome engineering strategy that enables flexible modeling of megabase-sized deletions. Generation of 9p21.3-syntenic deletions in a mouse model of pancreatic cancer revealed that concomitant loss of *Cdkn2a/b* and the IFN cluster led to immune evasion and metastasis compared to *Cdkn2a/b*-only deletions. Mechanistically, IFN co-deletion disrupted type I IFN signaling, altered antigen-presenting cells, and facilitated escape from CD8+ T cell surveillance in a cell extrinsic manner requiring loss of interferon epsilon (*Ifne*). Our results establish co-deletions of the IFN cluster as a pervasive route to tumor immune evasion and metastasis, revealing how deletions can disable physically linked cell intrinsic and extrinsic tumor suppression. Our study establishes a framework to dissect the functions of genomic deletions in cancer and beyond.

## MAIN

Understanding the genetic underpinnings of cancer is a fundamental goal of cancer research. Most efforts have focused on the characterization of single nucleotide variants (SNVs), which typically act as ON/OFF switches that affect the output of a single gene. An even larger class of cancer-associated lesions are copy number alterations (CNAs), which simultaneously impact the dosage of multiple genes and include chromosomal gains and losses, focal amplifications, and heterozygous or homozygous deletions^1,2^. Current estimates suggest that a typical tumor carries an average of 24 distinct CNAs that impact up to 30% of the genome^3,4,6^. Moreover, CNAs show recurrent patterns that can be associated with clinical outcomes^3,4,7,8^, arguing for active selection of specific traits rather than stochastic accumulation of genomic alterations. While much of the research on CNAs has focused on known drivers within the affected regions, emerging evidence indicates that co-gained or co-deleted genes – once considered “passenger” events – can also contribute to tumorigenesis^1,9,10^. While these observations imply CNAs produce complex phenotypes that cannot be recapitulated by manipulating a single gene^11-17^, the experimental modeling of these lesions remains a major challenge that has impeded the functional assessment of CNA biology^12,13,18-21^.

Among recurrent CNAs, loss of chromosome 9p21.3 is most strongly linked to poor prognosis and the most common homozygous deletion across human cancers^3,7^. The 9p21.3 locus is particularly prominent since it encompasses multiple key tumor suppressor genes (TSGs): the cell cycle inhibitors *CDKN2A* (encoding p16^INK4a^ and p14^ARF^) and *CDKN2B* (encoding p15^INK4b^), which collectively engage the function of p53 and RB, the major tumor-suppressive pathways that are impaired in cancer^5,22-24^. Hence, the current paradigm of how 9p21.3 deletions contribute to tumorigenesis is by eliminating a cell-intrinsic proliferative block. Nonetheless, several observations have been difficult to explain by this paradigm. Tumors with 9p21.3 deletions can display altered immune infiltrates^25,26^ and increased resistance to immune checkpoint blockade^27,28^, suggesting that the locus may also influence immune-related processes. Consistent with this possibility, numerous genome-wide association studies have identified single nucleotide polymorphisms in 9p21.3 even in non-cancer pathologies, notably age- and inflammation-related conditions^29^. However, the biological and molecular basis for these observations remains poorly understood.

### MACHETE Enables Efficient Generation of Megabase-sized Genomic Deletions in Cellular Models

To facilitate the experimental study of genomic deletions, we developed a rapid and flexible approach to engineer megabase-sized deletions termed <u>M</u>olecular <u>A</u>lteration of <u>Ch</u>romosomes with <u>E</u>ngineered <u>T</u>andem <u>E</u>lements (MACHETE). MACHETE involves an integrated process that inserts a selection cassette within a region of interest, followed by its co-deletion with defined regions of flanking DNA (**Figure 1A**). First, a bicistronic cassette encoding tandem negative and positive selection markers is amplified using oligos with homology to a region within an intended deletion. Second, the cassette is then inserted into the genome by CRISPR-facilitated homology-directed repair, and cells with integrations are enriched by positive selection. Third, a pair of single guide RNAs (sgRNAs) targeting the breakpoints of the intended deletion are introduced on either side of the bicistronic cassette, followed by negative selection. Since the sequence specificity of the flanking guides exclusively deletes on-target integrations of the suicide cassette, the latter step not only eliminates cells that retain the selection cassette but also those harboring off-target integrations (**Figure 1A**). Notably, the MACHETE protocol was designed to eliminate the need for cloning components: donor DNA is generated by introducing 40-bp homology arms via PCR amplification of the selection cassette, which is coupled to ribonucleoproteins (RNPs) of recombinant Cas9 complexed with sgRNAs (**Extended Data Figure 1A-B**). We envisioned that this approach would enable engineering of an allelic series of deletions, thereby enabling the systematic functional dissection of distinct regions within a locus.

**Figure 1.**
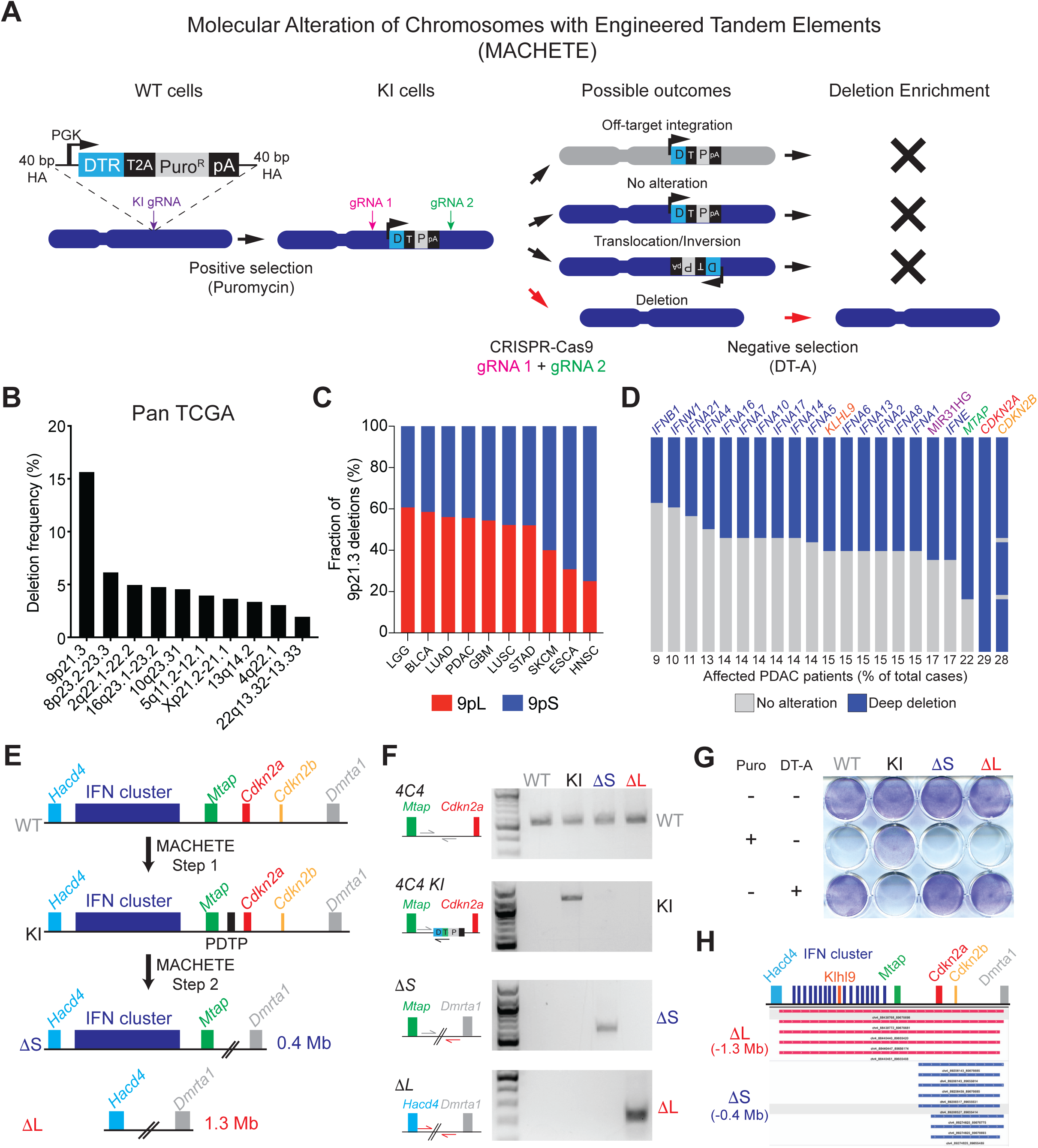
MACHETE Enables Efficient Engineering of Genomic Deletions. (A) Schematic of the MACHETE approach. (B) Frequency of homozygous deletions across the pan-cancer TCGA dataset. (C) Relative frequency of deletions at the 9p21.3 locus classified as 9pS and 9pL across different cancer types. (D) Frequency of deep deletion of 9p21.3 genes in PDAC patients. (E) Schematic of MACHETE-mediated engineering of 4C4 1′S and 1′L deletions. (F) PCR genotyping for the WT, KI, 1′S and 1′L alleles in the indicated PDEC cell lines. (G) Pattern of resistance/sensitivity to positive and negative selection in PDEC sgP53 EL parental, 4C4 KI, 1′S, and 1′L cells. Cells were seeded and treated with Puromycin (2 μg/mL) or DT-A (50 ng/mL) for 72 hours, and then stained with crystal violet to assess surviving cells. (H) DNA sequencing of breakpoints from 1′S and 1′L cells confirming loss of the expected genomic regions (0.4 Mb deletion in ΔS, and 1.3 Mb deletion in 1′L).

As an initial proof of concept, we engineered a 4.1-Mb deletion of the murine 11B3 locus (syntenic to human 17p13.1), which encompasses the *Trp53* TSG (**Extended Data Figure 1C**) and had been previously produced using a Cre/loxP approach^13^. NIH3T3 fibroblasts were targeted with a <u>P</u>GK-<u>D</u>TR-<u>T</u>2A-<u>P</u>uro (PDTP) dual-selection cassette to an intronic region of *Ccdc42*, a gene located in the 11B3 locus, and positively selected for insertion of the cassette (11B3 knock-in (KI) cells). Cas9-sgRNA RNPs were then introduced to target regions flanking *Sco1* and *Alox12,* the genes that demarcate the intended deletion, and negative selection was performed using DT to produce a cell population termed Δ11B3 (**Extended Data Figure 1C**). Parental, 11B3 KI, and Δ11B3 populations showed the expected pattern of resistance or sensitivity to the selection agents, presence/absence of the cassette, and expected deletion breakpoint (**Extended Data Figure 1D, E**). Clonal analysis showed that use of negative selection effectively enabled the generation of the desired deletion, by increasing the efficiency of Δ11B3 engineering from undetectable (0/22) to 40% of positive clones (11/27, all heterozygous) (**Extended Figure 1F**), which was confirmed by sequencing (**Extended Data Figure 1G**).

We further developed a series of constructs that enable the use of MACHETE across various experimental contexts (**Extended Data Figure 1H**). To illustrate the use of MACHETE in human cells, we selected a cassette composed by a herpes simplex virus thymidine kinase with blue fluorescent protein (HSV-TK-T2A-BFP), which enables positive selection via fluorescence activated cell sorting (FACS) and negative selection using ganciclovir. This construct enabled the production of cells harboring a 45 Mb deletion of chromosome 7q11-7q22 (**Extended Figure 1I-K**). Thus, MACHETE is a customizable approach to efficiently engineer large chromosomal deletion events.

### Loss of Type I IFN Genes Is a Common Event in 9p21.3 Deleted Tumors

Armed with MACHETE, we set out to interrogate the biology of deletions at the 9p21.3 locus (**Figure 1B**). Interestingly, although *CDKN2A* is a well-established tumor suppressor in this region, we and others have noted that 9p21.3 deletions can encompass additional genes, including a cluster of 16 type I IFN genes whose genetic loss has not been functionally implicated in tumorigenesis despite the known role of IFN signaling in anti-tumor immunity^30^. An analysis of the TCGA dataset^31^ revealed that fourteen different tumor types harbor homozygous 9p21.3 deletions in over 10% of cases (**Extended Data Figure 2A**). We further classified 9p21.3 deletions into those targeting *CDKN2A/B* alone (9p small, or 9pS) or larger events that typically encompassed the entire type I IFN cluster (9p large, or 9pL) (**Figure 1C**). The frequency of the 9pL events ranged between 20-60% depending on tumor type and was one of the highest in pancreatic ductal adenocarcinoma (PDAC) (**Figure 1D**).

### Engineering 9p21.3 Deletions in Mouse Models of PDAC

Genetic analyses of human PDAC indicate that *CDKN2A* deletions are an early event in tumor evolution^32,33^, which are thought to emerge as heterozygous deletions that subsequently undergo loss of heterozygosity^34,35^. These deletions tend to co-occur with activating *KRAS* mutations and *TP53* loss, two other major drivers in this disease (**Extended Data Figure 2B**)^36^. Given the potential role of type I IFNs in modulating immune responses, we set out to study the biology of different 9p deletions in a syngeneic model of murine PDAC derived from established pancreatic ductal epithelial cells (PDECs) that harbor an endogenous activated Kras^G12D^ allele^37,38^. While *Cdkn2a* expression is blunted in this system, the lesions produced following PDEC transplantation resemble premalignant stages of PDAC, display a limited capacity to progress to invasive adenocarcinoma^38^, and allow the study of immune-related processes^37,39^. Thus, given the synteny between human 9p21.3 and murine 4C4 (**Extended Data Figure 2C**), PDEC cells provide a good platform for MACHETE-based engineering of 9p21.3 equivalent deletions *in vitro* and the subsequent study of tumor phenotypes in an immune competent in vivo context.

To model the most relevant genetic configuration for 9p21.3 loss in human PDAC, we generated *Trp53* knockout PDEC cells using transient CRISPR-Cas9 and introduced an EGFP-Luciferase cassette to enable visualization of engrafted cells (PDEC-sgP53-EL cells) (**Extended Data Figure 2D**). MACHETE was then used to engineer the two most frequent configurations of 9p21.3 deletions: ΔS (“Small”; 0.4 Mb loss spanning *Cdkn2a* and *Cdkn2b*), and ΔL (“Large”; 1.3 Mb loss spanning the entire 4C4 locus) (**Figure 1E-G**). Deep sequencing of the breakpoint regions confirmed the presence of precise 0.4 and 1.3 Mb deletions, and clonal analysis of targeted cell populations indicated that MACHETE achieved an >8-fold increase in producing cells with the intended heterozygous deletion (**Figure 1H**, **Extended Data Figure 2E, F**). As expected, these populations could be further edited through MACHETE’s capability for iterative engineering (**Extended Data Figure 2G**). Given the comparable deletion efficiency of ΔS and ΔL cells, cell populations were used for subsequent analyses to minimize the effects of clonal variation.

### Tumors with ΔL Deletions Are Differentially Surveilled by the Adaptive Immune System

To determine whether each heterozygous deletion event contributes to tumorigenesis, we transplanted the ΔS and ΔL lines into the pancreata of syngeneic C57BL/6 recipients and assessed tumor formation at 4 weeks via bioluminescent imaging and at endpoint. Cells bearing the ΔL deletion tended to form more tumors than ΔS cells, although the difference was not statistically significant (**Figure 2A**). Tumors arising from both genotypes were poorly differentiated, consistent with the histopathology of autochthonous *Trp53-* and *Cdkn2a*-deficient PDAC models (**Extended Data Figure 2H**)^40^. Sparse whole genome sequencing (sWGS) confirmed that most ΔS or ΔL tumors acquired homozygous deletions of their respective alleles (7/9 lines for ΔS; 6/8 lines for ΔL), as occurs in human PDAC (**Extended Data Figure 2I**). However, there was one notable difference: ΔL tumors retained a strong EGFP fluorescence signal and genomic copy number compared to ΔS tumors (**Figure 2B-C**).

**Figure 2.**
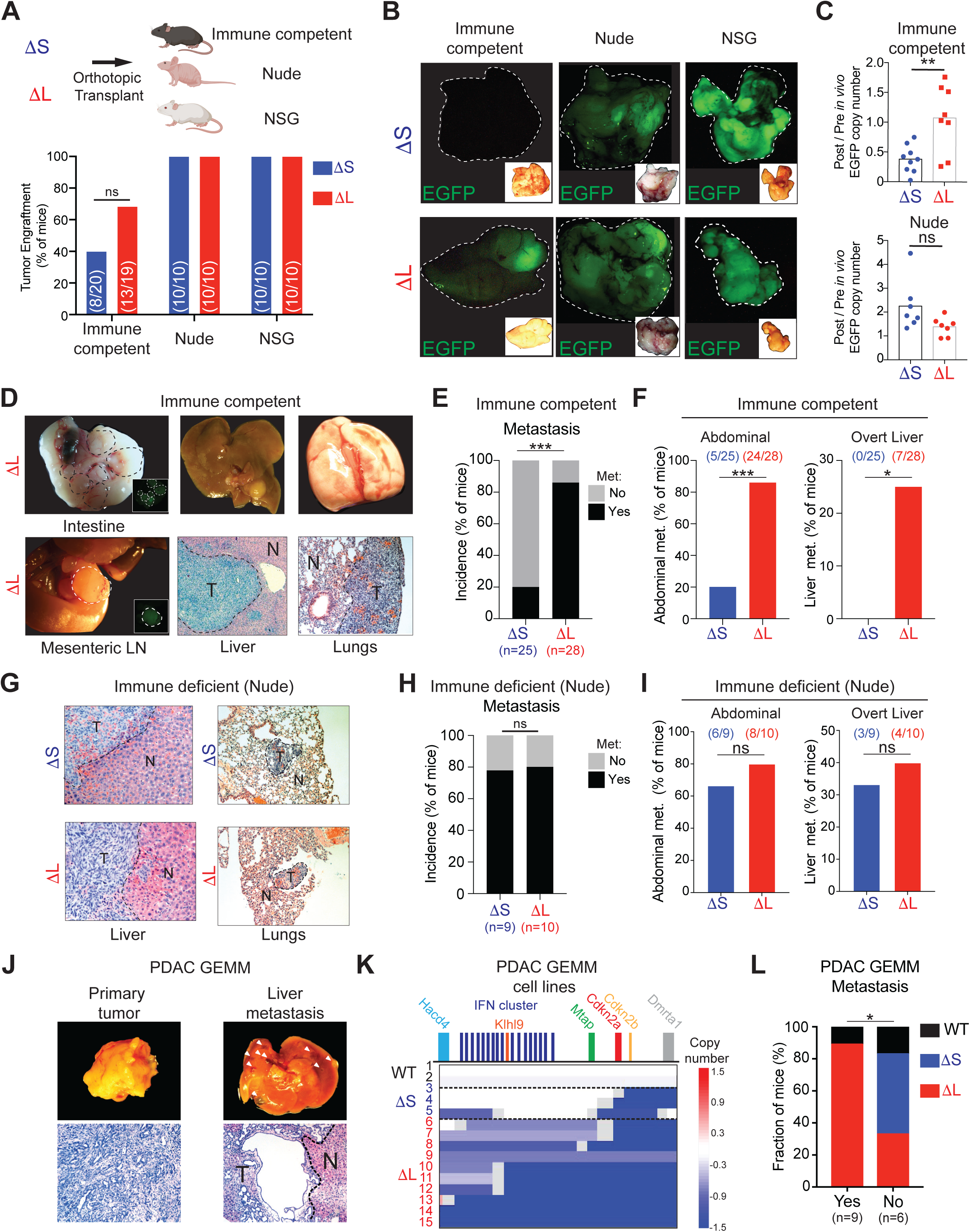
1′L Deletions Are Differentially Surveilled by the Adaptive Immune System and Promote Metastasis. (A) Engraftment at one month after injection of 1′S and 1′L cells in C57BL/6, nude, and NSG hosts. Two independently generated input cell lines were used per genotype (n ζ 5 per each cell line). Bars represent fraction of metastasis-bearing mice (specific numbers of independently analyzed mice are noted in parentheses). ns = non-significant (chi-square test). (B) Representative macroscopic fluorescent images of primary tumors harvested from the indicated genotypes and hosts. Insets show the brightfield image for each tumor. (C) qPCR analysis for EGFP copy number in the gDNA of tumor-derived (Post *in vivo*) 1′S and 1′L lines from C57BL/6 and Nude hosts, relative to their parental (Pre *in vivo*) counterparts. Each dot represents an independent tumor-derived cell line. **p < 0.01, ns = non-significant, two-tailed t-test. (D) Representative images of metastases in C57BL/6 mice with 1′L tumors. Left: Brightfield macroscopic images of abdominal (intestinal and mesenteric lymph node) metastases. Insets show matched EGFP fluorescence images. Middle: Macroscopic and Hematoxylin/Eosin images of tumor-bearing livers. Right: Macroscopic and Hematoxylin/Eosin images of tumor-bearing lungs. T = tumor; N = normal adjacent tissue. (E-F) Overall (E) and organ-specific (F) metastasis incidence in C57BL/6 mice with either 1′S or 1′L tumors. 4 independently generated input cell lines were used per genotype (n ζ 5 per each cell line). Bars represent fraction of metastasis-bearing mice (specific numbers of independently analyzed mice are noted in parentheses). *p < 0.05; ***p < 0.001, chi-square test. (G) Representative images of metastases in Nude mice with 1′L or 1′S tumors. Hematoxylin/ Eosin images of tumor-bearing livers (left) and lungs (right) are shown. (H-I) Overall (H) and organ-specific (I) metastasis incidence in C57BL/6 mice with either 1′S or 1′L tumors. 4 independently generated input cell lines were used per genotype (n Δ 5 per each cell line). Bars represent fraction of metastasis-bearing mice (specific numbers of independently analyzed mice are noted in parentheses). ns = non-significant, chi-square test. (J) Representative gross morphology (top) and Hematoxylin/Eosin histological stain (bottom) of matched primary tumor and overt liver metastasis in a Kras^G12D/+^; shSmad4 PDAC GEMM. (K) sWGS analysis of tumor-derived cell lines from the KC-Ren and KC-Smad4 GEMMs, grouped by spontaneous 4C4 deletion type (WT, 1′S, 1′L). Schematic of the murine 4C4 locus is shown on top. Blue tracks indicate deleted regions, with color intensity corresponding to the extent of the deletion. Numbers correspond to independent mice. (L) Incidence of overt metastasis in mice with tumors that harbor WT 4C4 locus, 1′S or 1′L deletions. Bars represent fraction of metastasis-bearing mice (specific numbers of independently analyzed mice are noted in parentheses). *p < 0.05, chi-square test.

The above findings are consistent with immunoediting of cells with high reporter expression^41^ and raise the possibility that ΔL cells may be less immunogenic than their ΔS counterparts. Accordingly, ΔS and ΔL cells showed a similar capability of forming EGFP-expressing tumors in *Foxn1^nu^*(“nude”: T and B cell deficient) and NOD/SCID *Il2rg^-/-^* (NSG: T, B, and NK cell deficient) mice (**Figure 2A-C**). Interestingly, cell populations engineered to harbor 4C4 deletions that eliminate upstream elements but that retained the *Cdkn2a/b* genes (ΔI allele) had reduced tumor initiating capacity yet produced tumors that expressed similar levels of EGFP as ΔL tumors (**Extended Data Figure 2J-L**). These data imply that genetic elements upstream of *Cdkn2a/b* contribute to immunoediting of developing tumors.

### ΔL Deletions Promote Metastasis by Evasion of Adaptive Immunity

We next compared the behavior of ΔS and ΔL tumor-derived cell lines in orthotopic transplantation assays. Four independently derived ΔS and ΔL tumor lines were FACS-sorted to obtain cell populations with comparable EGFP levels to eliminate differences in reporter expression as a confounding factor (**Extended Data Figure 3A**). ΔS and ΔL tumor cells showed a similar ability to proliferate in adherent or suspension cultures and produced tumors with undifferentiated histopathology (**Extended Data Figure 3B, C**). However, consistent with their acquisition of homozygous 4C4 deletions, the tumors progressed more rapidly (**Extended Data Figure 3D**).

Although ΔS and ΔL tumors showed no obvious difference in the fraction of proliferating or apoptotic cells (**Extended Data Figure 3E**), ΔL tumors were much more prone to metastasis (**Figure 2D-E**). Indeed, these mice displayed a 4-fold higher incidence of macrometastases in the abdominal area (mesenteric lymph nodes, intestine, and peritoneal cavity) compared to their ΔS counterparts, and uniquely harbored overt liver metastases (∼25% of mice) (**Figure 2F**). These observations were confirmed through histological analyses, which also indicated a trend for larger number and area of liver lesions (**Extended Data Figure 3F-G**).

Further insights into the greater metastatic potential of ΔL cells were obtained through analyzing additional tumor genotypes and routes of cell delivery, or by studying tumor behavior in immunocompromised animals. First, tumor-derived cells that remained heterozygous for the ΔL (2/8 lines that did not undergo LOH) or ΔI alleles were unable to efficiently produce metastases following orthotopic injection (**Extended Data Figure 3H-I**). Second, homozygous ΔS or ΔL tumor cells were equally able to produce experimental liver metastases following intrasplenic injection (**Extended Data Figure 3J-K**). Third, as was observed for the immunoediting phenotype, homozygous ΔS and ΔL cells showed a similarly high frequency of metastasis following orthotopic injection into Nude mice (**Figure 2G-I**, **Extended Data Figure 3L-M**). Therefore, the enhanced metastatic propensity of ΔL cells requires concomitant homozygous deletion of *Cdkn2a/b* and the IFN cluster and involves an immune surveillance mechanism that acts prior to the colonization at distant sites.

### Loss of Type I IFNs Correlates with Metastasis in Autochthonous Mouse Models of PDAC

Next, we tested the association between large 4C4 deletions and metastasis in an independent and autochthonous genetically engineered mouse model (GEMM) of PDAC. In agreement with a previous report^36^, metastatic GEMM tumors initiated by mutant *Kras^G12D^*alone or in combination with a TGFβ pathway alteration (*Smad4* depletion in our model) spontaneously acquire 4C4 deletions during their natural course of tumor evolution (**Figure 2J-L**). Analysis of deletion size revealed that primary tumor cells isolated from mice with metastases almost always harbored large 4C4 events (8/9 mice) whereas those without overt metastases had focal *Cdkn2a/b* deletions or no 4C4 alteration (4/6 mice) (**Figure 2K, L**). The presence and extent of 4C4 deletion was similar between individual primary and metastatic pairs (n=7), confirming that 4C4 loss is an early event in this model (**Extended Data Figure 3N**). Nonetheless, in contrast to the PDEC system, primary tumors arising in this GEMM model displayed a moderately differentiated histology with stromal involvement (**Figure 2J**), implying that the increased metastatic potential associated with large 4C4 deletions does not require an undifferentiated histopathology. These orthogonal data reinforce the notion that one or more genes unique to the 1′L deletion suppress metastasis.

### 4C4 Deletion Genotype Dictates Type I IFN Signaling and Immune Infiltration

To help understand how distinct 4C4 deletion events influence tumor phenotypes, we performed RNA-seq on bulk 1′L and 1′S tumors and inferred differences in signaling pathways and immune cell composition using CIBERSORT^42^. When compared to 1′S tumors, 1′L tumors displayed a decrease in pathways linked to IFN signaling (**Extended Data Figure 4A-B**), as well as a broad depletion in immune signatures, including B and T cell populations (**Extended Figure 4C**). Further analyses using RT-qPCR confirmed that 1′L tumors have reduced levels of type I IFNs (*Ifnb1* and *Ifne*) and IFN-responsive genes (*Oasl1* and *Isg20*) (**Extended Data Figure 4D**). Adding granularity to these observations, single cell RNA sequencing (scRNA-seq) of tumor-infiltrating CD45+ cells isolated from 1′S and 1′L tumors identified changes in the abundance of multiple immune cell populations (**Extended Data Figure 4E-I**). 1′L tumors had fewer B cells and myeloid populations, which was accompanied by an increase in CD8+ T cells – changes that were confirmed by flow cytometry (**Figure 3A-B**, **Extended Data Figure 5A-I**).

**Figure 3.**
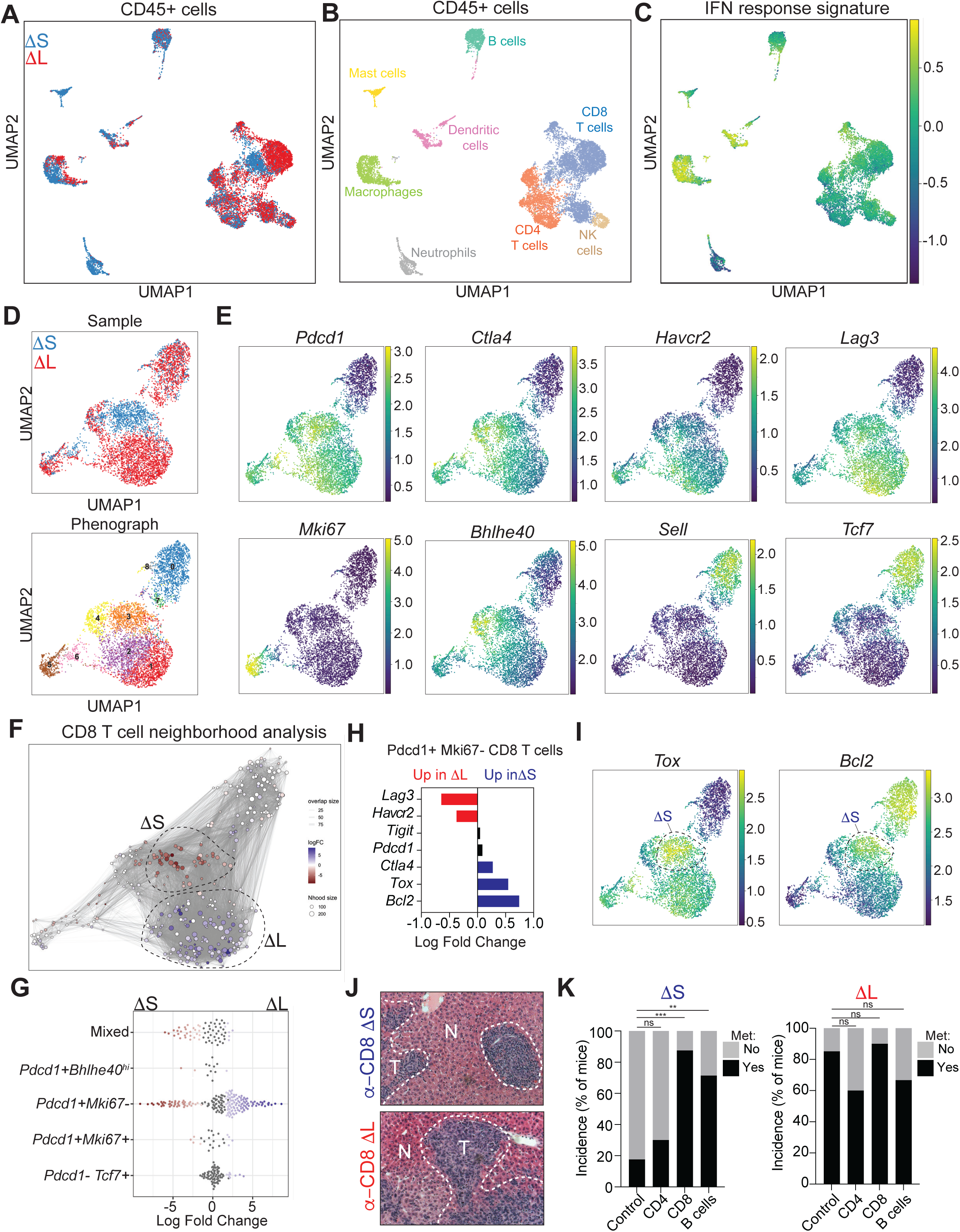
4C4/9p21.3 Deletion Genotype Dictates Type I IFN Signaling and Immune Infiltration. (A) UMAP of CD45+ cells showing cells derived from 1′S (n = 7774 cells) or 1′L (n = 7560 cells) tumors. (B) UMAP of CD45+ cells annotating the specific immune subsets. (C) UMAP of averaged IFN response signature across CD45+ populations. (D) (Upper) UMAP of CD8+ T cells from 1′S or 1′L tumors. Cells are colored by sample. (Bottom) UMAP of CD8+ T cell clusters. Cells are colored and by their cluster identity. (E) UMAP of imputed expression for the indicated genes. (F) MILO analysis of CD8+ T cells. Neighborhoods identified through MILO analysis using default parameters (red indicates enrichment in 1′S, while blue indicates enrichment in 1′L). (G) Swarm plot of the distribution of CD8+ T cell neighborhoods in 1′S or 1′L tumors across transcriptional states. The x-axis indicates the Log-fold change in differential abundance of 1′S (<0) and 1′L (>0). Each neighborhood is associated with a cell type if more than 80% of the cell state in the neighborhood belong to said state, else is annotated as “Mixed”. (H) Differential gene expression of the indicated genes in *Pdcd1+ Mki67-* CD8+ T cells. (I) UMAP of imputed expression of *Tox* and *Bcl2*. Dashed circles highlight 1′S-enriched CD8+ T cells. (J) Representative images of liver metastasis upon CD8+ cell depletion. (K) Incidence of metastasis upon depletion of immune subsets in 1′S or 1′L tumors. 2 independently generated input cell lines were used per genotype (n Δ 5 per each cell line). Bars represent fraction of metastasis-bearing mice (specific numbers of independently analyzed mice are noted in parentheses). ns = non-significant; **p < 0.01; ***p < 0.001, chi-square test.

Beyond alterations in the composition of infiltrating CD45+ cells, the distinct 4C4 deletions led to changes in the transcriptional state of immune subsets. Analysis of an experimentally derived type I IFN response signature (see Methods and Extended Data Table 1) showed that professional antigen-presenting cells (APCs; macrophages, dendritic cells and B cells) and CD8+ T cells exhibited reduced type I IFN signaling in the 1′L setting (**Figure 3B-C**, **Extended Data Figure 4J**). Moreover, the specific effects of 4C4 deletions on APCs were immune cell type-dependent: a more pro-inflammatory state of cDC2 dendritic cells in ΔS tumors (**Extended Data Figure 5J-L**); a shift in macrophage transcriptional states toward higher M1-like cells in in 1′S tumors (**Extended Data Figure 5M-O**); and an overall reduction across all B cell subtypes in 1′L tumors (**Extended Data Figure 5P-Q**).

Analysis of CD8+ T cells showed a range of activation states, with a dominant presence of activated/exhausted (*Pdcd1*+, *Ctla4*+, *Havcr2*+, *Lag3*+), naïve (*Pdcd1*-, *Tcf7*+, *Sell*+), and cycling cells (*Pdcd1*+, *MKi67*+) (**Figure 3D-E**). Intriguingly, the non-proliferating *Pdcd1*+ population of CD8+ T cells occupied distinct phenotypic space in 1′S and 1′L tumors. Further characterization using MILO^43^ revealed that ΔS tumors accumulated exhausted CD8+ T cells marked by *Tox* and *Bcl2* expression, whereas those present in ΔL tumors were transcriptionally distinct and displayed higher expression of *Havcr2* and *Lag3* (**Figure 3F-I**, **Extended Data Figure 5R**, Extended Data Table 1). The high levels of IFN-engaged APCs and distinct CD8+ T cell states present in ΔS tumors implied ongoing immune surveillance that, based our phenotypic data, may suppress metastatic spread. In agreement, depletion of B and CD8+ cells, but not CD4+ cells, enhanced the metastatic potential of 1′S tumor cells to levels observed for 1′L tumors (**Figure 3J-K**). Collectively, these data suggest that loss of tumor-intrinsic type I IFNs impairs the function of professional APCs and produces a unique state of CD8+ T cell dysfunction, leading to defects in anti-tumor immunity.

### 9p21.3 Deletions Correlate with IFN signaling and Immune Infiltration in Human PDAC

To test how 9p21.3 deletions that encompass the type I IFN cluster alter the tumor microenvironment in human PDAC, we analyzed sequencing data obtained from the COMPASS trial, which contains 218 primary and 180 metastatic PDAC samples isolated by laser capture microdis section ^44,45^. The availability of whole genome and RNA sequencing from each of these samples allows tumors to be categorized based on 9p deletion status and then analyzed for immune signatures linked to infiltrating stromal cells. Consistent with our findings in murine tumors, analysis of primary tumors showed that 9pL deletions correlated with reduced type I IFN signaling compared to their 9pS counterparts (**Extended Data Figure 6A**).

The genotype-specific differences in gene ontology pathways and inferred immune cell composition correlated well across species (**Extended Data Figure 6B-C**, Extended Data Table 2). Notably, IFN cluster-proficient (ΔS/9pS) tumors were enriched in pathways associated with immune infiltration of both innate and adaptive categories (**Extended Data Figure 6B**) and showed a relative enrichment of most immune cell populations, particularly effector CD8+ T and B cell subsets (**Extended Data Figure 6C**). Nevertheless, the relative enrichment in type I IFN signatures present in primary 9pS tumors was reduced in 9pS metastases (**Extended Data Figure 6D**)^46^, and analysis of RNA-seq data from a second cohort of matched primary and metastatic PDAC samples confirmed a reduction in type I IFN signaling in metastases irrespective of tumor genotype (**Extended Data Figure 6E**). When considered in the context of our functional studies, these data imply that downregulation of type I IFN signaling, by genetic or other means, promotes PDAC metastasis.

### Disruption of IFNAR Signaling Phenocopies the Immune Evasive and Pro-metastatic Properties of **1′**L Cells

Besides type I IFNs, 1′L deletions include other genes, including *Mtap*, whose disruption can also influence tumor cell behavior^47^. To specifically test whether type I IFN signaling is required for the immune evasive and pro-metastatic features of 1′L tumors, we used IFNAR1 blocking antibodies as an orthogonal approach to disrupting type I IFN signaling in the host. Immune competent mice were pre-treated with an IFNAR1-blocking antibody or an isotype control, followed by orthotopic transplantation of 1′S and 1′L cells analysis of the resulting tumors for immunoediting of the EGFP-Luciferase reporter and overall incidence of metastasis (**Extended Data Figure 7A**).

Consistent with our model, 1′S tumors arising in mice subjected to IFNAR1 blockade expressed higher levels of EGFP than isotype-treated controls (**Figure 4A**, **Extended Data Figure 7B-C**) and showed a greater incidence of metastasis in secondary transplantation assays (**Figure 4B**; **Extended Data Figure 7D-F**). Remarkably, these patterns were comparable to those arising in immune competent mice receiving ΔL cells and in immune deficient animals transplanted with ΔS cells (**Figure 2C, F, I**). In contrast, type I IFN blockade had no impact on the enhanced metastatic potential of 1′L cells (**Figure 4B**). Transcriptional profiling of bulk tumors confirmed that IFNAR1 blockade phenocopied the reduction of type I IFN signaling observed in IFN-deficient tumors but had minimal impact on the transcriptome of 1′L tumors (**Figure 4C**; **Extended Data Figure 7G-H**). These data imply that that one or more type I IFNs are required for the immune evasive and pro-metastatic phenotypes arising in tumors with homozygous 1′L deletions.

**Figure 4.**
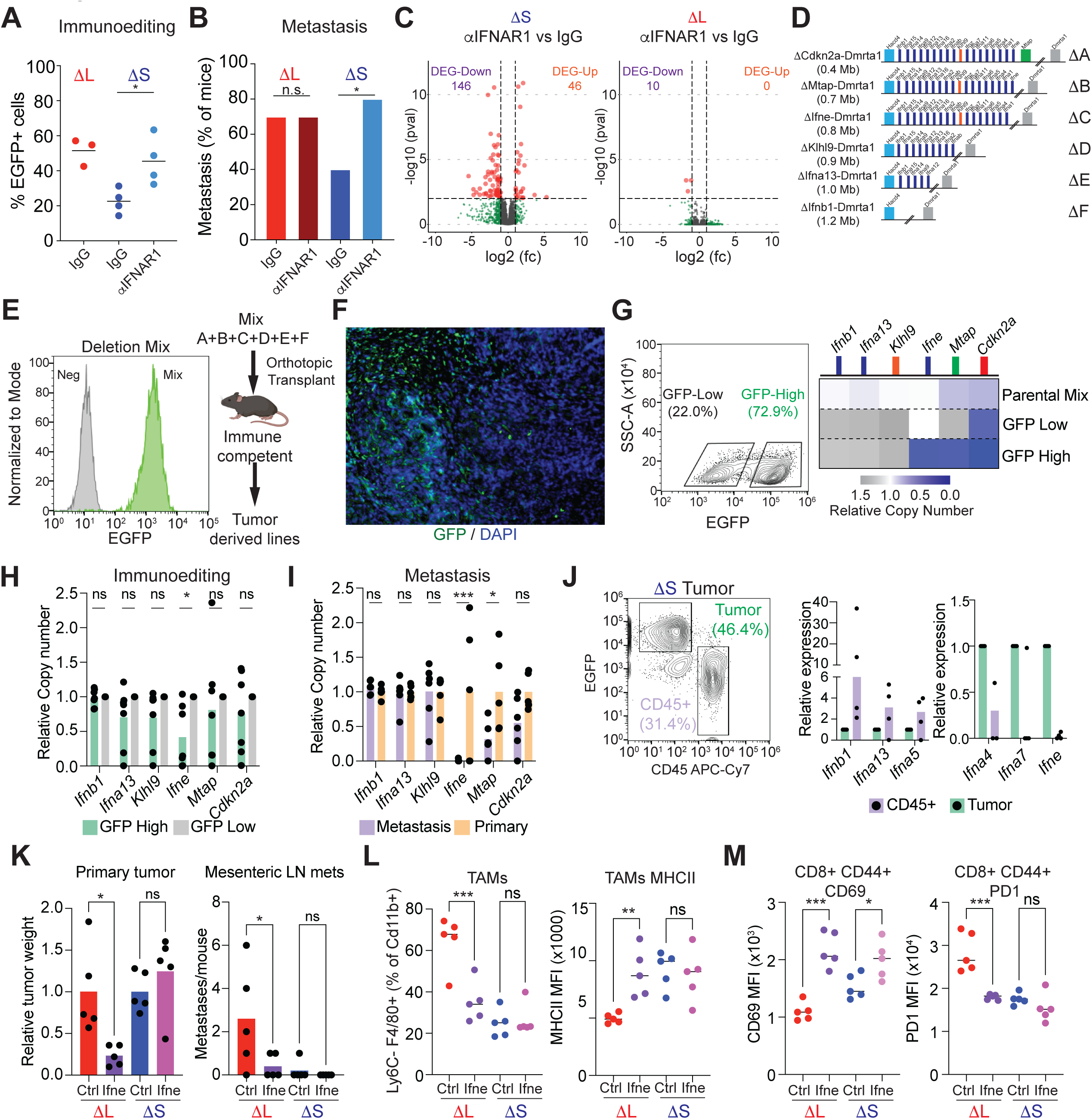
*Ifne* Is a Tumor-specific Mediator of Immune Surveillance and Metastasis. (A) Quantification of EGFP fluorescence in 1′S or 1′L tumors from C57BL/6 mice treated with IgG or αIFNAR1 antibodies. Representative plots are shown in **Extended Data Figure 7B**. Each dot represents an independent biological replicate. *p < 0.05, one-way ANOVA followed by Tukey’s multiple comparison test. (B) Incidence of metastasis in C57BL/6 mice transplanted with homozygous 1′S or 1′L lines and treated with IgG or αIFNAR1 antibodies. 2 independently generated input cell lines were used per genotype (n Δ 5 per each cell line). Bars represent fraction of mice bearing metastasis (total numbers of independently analyzed mice are shown). ns = non-significant; *p < 0.05, chi-square test. (C) Volcano plots of differentially expressed genes comparing IFNAR1 blockade vs. IgG controls in 1′S or 1′L tumors. (D) Schematic of extended series of 4C4 deletion alleles. (E) (Left) Flow cytometry measurement of EGFP fluorescence in tumors derived from deletion series mix (“Mix”). EGFP-negative cells were used as negative controls (“Neg”). (Right) Schematic of *in vivo* competition experiment. (F) Representative EGFP immunofluorescent stain of a deletion-mix tumor. (G) (Left) Representative flow cytometry plot of EGFP levels in a deletion-mix tumor. GFP-Low and GFP-High cell populations were sorted as marked. (Right) Copy-number qPCR analysis of the indicated genes in the parental cell mix, and GFP-Low vs. GFP-High cells sorted from resulting tumors. (H) Relative copy-number quantification of indicated genes in GFP-High vs. GFP-Low cells. *p<0.05; ns=non-significant, one-way ANOVA followed by Sidak’s multiple comparison test. Bars represent SEM, n=7 biological replicates. (I) Relative copy-number quantification of indicated genes in metastases-vs. primary tumor-derived cells. *p<0.05; ***p<0.001; ns=non-significant, one-way ANOVA followed by Sidak’s multiple comparison test. Bars represent SEM, n=7 primary tumors and 6 metastases. (J) RT-qPCR measurements of mRNA levels for the indicated IFN genes in tumor cells and infiltrating CD45+ cells from 1′S tumors. Each dot is a biological replicate (n=4). (K) Relative quantification of primary tumor weights (left) and number of mesenteric LN metastases (right) in 1′S and 1′L tumors with add-back of Ifne-expressing or control construct. *p<0.05; ns=non-significant, one-way ANOVA followed by Sidak’s multiple comparison test to the respective control population. Each dot is an independent tumor. (L) Flow cytometry-based quantification of TAM fraction (left) and TAM MHCII levels (right) in tumors of the indicated genotypes. **p<0.01; ***p<0.001; ns=non-significant, one-way ANOVA followed by Sidak’s multiple comparison test to the respective control population. Each dot is an independent tumor. (M) Flow cytometry-based quantification of CD69 (left) and PD1 (right) levels in CD8+CD44+ T cells from tumors of the indicated genotypes. *p<0.05; ***p<0.001; ns=non-significant, one-way ANOVA followed by Sidak’s multiple comparison test to the respective control population. Each dot is an independent tumor.

### *Ifne* Is a Tumor-specific Mediator of Immune Surveillance and Metastasis

The functional redundancy between different type I IFNs remains poorly understood^48^. For instance, *Ifnb1* is highly expressed in immune cells and acts as a key downstream effector of the cGAS-STING pathway to engage innate and adaptive immunity, yet the individual contributions of most other IFNs to infection and cancer immunity are unclear^30,49^. To dissect the functional contribution of tumor-derived IFNs to immunoediting and metastasis, we leveraged the power of MACHETE to engineer a refined deletion series that encompass a gradually increasing number of IFN genes (**Figure 4D**). The resulting cell populations were orthotopically injected as pools into immunocompetent recipient mice (**Figure 4E**) and expression of EGFP-Luc reporter was used as an indicator of immune evasion in the resulting tumors.

Consistent with different deletion events affording different degrees of immune evasion, the tumors showed heterogenous expression of EGFP (**Figure 4F**). Isolation of cells with distinct levels of EGFP showed prevalence in deletions affecting the IFN cluster in the EGFP-retaining population (**Figure 4G**), with a significant enrichment of cells harboring deletions of *Ifne* across multiple independent tumors (**Figure 4H**). A similar increase in the deletion of *Ifne* was observed when comparing metastases to primary tumors, further highlighting the potential relevance of *Ifne* to tumor dissemination (**Figure 4I**).

A detailed analysis of type I IFN gene expression in epithelial and CD45+ immune cells present in ΔS tumors reinforced the above observations. As previously reported, *Ifnb1* could be induced by a cGAS-STING agonist yet was more highly expressed in immune cells than tumor cells; by contrast, other IFNs, particularly *Ifne*, were not induced by these stimuli and showed preferential expression in tumor cells (**Figure 4J**, **Extended Data Figure 7I-J**). Collectively these data imply that disruption of *Ifne* is necessary for the effects of type I IFN loss on immune evasion and metastasis.

To determine whether *Ifne* was sufficient to suppress immune evasion and metastasis, we introduced a doxycycline-inducible construct to drive either full-length *Ifne* or a truncated *Ifne* control in 1′S and 1′L cells (**Extended Data Figure 8A-C**). Both sustained and acute induction of full-length *Ifne* suppressed overt metastasis of 1′L tumors, which was dependent on adaptive immunity (**Figure 4K**, **Extended data Figure 8D-H**). Despite the expected overexpression of *Ifne* and downstream type I IFN target genes (**Extended Data Figure 8I**), 1′S and 1′L tumors showed differential response to acute *Ifne*: 1′S tumors had no effect on primary tumor growth while 1′L tumors had a reduction in tumor size and metastasis (**Figure 4K**). Consistent with loss of function phenotypes, tumors with enforced *Ifne* expression displayed elevated levels of professional antigen-presenting populations and an increase in activated CD8+ T cells (**Figure 4L-M**, **Extended Data Figure 8J**). Taken together, these data demonstrate that somatic deletion of type I IFNs impairs immunoediting and metastasis via the adaptive immune system and reveal a previously unanticipated role of *Ifne* in suppressing these phenotypes.

## DISCUSSION

Despite the pervasive nature of CNAs across cancers, their functional characterization has been hampered by the difficulty of manipulating large genomic regions. MACHETE addresses this challenge by providing an efficient method that is customizable to any genomic locus, enables the engineering of deletions of at least 45 Mb in size, and is easy to adopt: it requires no cloning of targeting vectors, seamlessly eliminates cells with off-target integrations, and – as shown herein – allows for iterative engineering of refined deletions or increasingly complex genotypes. Using MACHETE, we reveal previously unappreciated but clinically relevant insights into the multifactorial nature of 9p21.3 deletions, an event that contributes to up to 15% of human cancers^50^. Given the emerging view that CNAs influence cancer phenotypes by altering the dosage of multiple genes, tools like MACHETE will be essential for understanding their biology and any therapeutic opportunities they create.

Our results revise the long-standing paradigm for how genes encoded at the 9p21.3 locus suppress tumorigenesis. Most studies have focused on the roles of the *CDKN2A* (encoding p16^INK4a^ and p14^ARF^), and to a lesser extent, *CDKN2B* genes (encoding p15^INK4b^), which act in concert to potently suppress tumorigenesis by driving premalignant cells into a stable state of cell cycle arrest^5^. Herein, we show that the type I IFN cluster is co-deleted with *CDKNA/B* in nearly half of all tumors harboring 9p21.3 deletions and encodes factors that act as potent tumor-derived enforcers of anti-tumor immunity. While other 9p21.3 genes such as *Mtap^47,51^* may also influence tumors, our data pinpoint type I IFNs as the phenotypically important tumor suppressors in our model. Therefore, 9p21.3 deletions not only disable a potent block to cancer proliferation but also facilitate immune evasion, simultaneously disrupting cell intrinsic and extrinsic tumor-suppressive programs.

The emerging picture from our data argues that *Cdkn2a* loss is a requisite event that enhances proliferate capacity while co-deletion of type I IFNs provides a collateral benefit that promotes immune evasion by altering immune infiltrates in the developing tumor. This model also explains why neighboring cells are unable to compensate for type I IFN deletions, as incipient tumors may eventually reach a size where paracrine IFN signaling becomes limiting. Regardless, the ability of tumor cells harboring type I IFN deletions to avoid immune surveillance at the primary tumor site increases their metastatic potential. As such, the type I IFN cluster acts as a bona fide metastasis suppressor locus, adding support to the emerging view that immune surveillance plays an important role in limiting metastatic spread and contrasting with the prevailing model that metastasis is strictly driven by epigenetic changes.

The role of different tumor-derived type I IFNs during cancer progression has remained unclear, with most attention given to IFN secretion by immune cells or the regulation of *Ifna/b* genes downstream of cGAS-STING signaling^49,52-55^. Nonetheless, in our model a subset of type I IFNs, particularly *Ifne*, are exclusively expressed in tumor cells, where they promote type I IFN signaling and dictate the composition and state of immune cell infiltrates. Consequently, deletion of the type I IFN cluster produces a tumor microenvironment that culminates with the accumulation of exhausted CD8+ T cells that display markers of terminal differentiation, analogous to those observed in IFNAR1 knockout mice during defective responses to pathogen challenge^56^. The lack of induction of *Ifne* in response to classic type I IFN inducers (such as TLR and cGAS-STING agonists) highlights its unique role as a potentially constitutive enforcer of tumor immune surveillance, perhaps mirroring its only known role in mediating mucosal immunity^57^.

In sum, our results nominate type I IFN deletions as a pervasive genetic mechanism of immune evasion in cancer, rivaling heterozygous deletions of the HLA cluster^58^, and as such may explain the correlation between 9p21.3 deletions and resistance to immune checkpoint blockade^50,59,60^. Whether the physical coupling between IFNs and *Cdkn2a/b* is biologically meaningful or coincidental remains to be determined, but it is noteworthy that both type I IFNs and *Cdkn2a*-encoded proteins have roles in limiting viral infection^48,61^ that may have been co-opted for tumor suppression. Intriguingly, genome-wide association studies have identified the 9p21.3 locus as one of two highly significant regions that are broadly associated with a series of age-related pathologies, the other key region remarkably coinciding with the HLA locus on chromosome 6p21^29,62^. While *CDKN2A* is thought to drive the 9p associations, the cooperative effects between *CDKN2A* and type I IFNs reported herein raise the possibility that variation in type I IFN regulation plays a role in the biology of these pathologies as well.

## METHODS

### Pan-cancer TCGA Data Analysis

Analysis of TCGA datasets was performed on cBioPortal^63,64^. All TCGA datasets were selected and the following onco-query language (OQL) entry was used (Extended Data Table 3 for 9p21.3 OQL). Tumors with at least 10% of patients harboring 9p21.3 deletion were identified. Tumors were classified as 9pS if they had a focal deep deletion of *CDKN2A/B*. Tumors were classified as 9pL if both *CDKN2A/B* and the type I IFN cluster was deleted. For the 9pL/9pS relative frequency, only datasets with at least 40 cases with 9p21.3 loss were considered.

### Cell Culture

NIH3T3 fibroblasts were obtained from the American Type Culture Collection (ATCC), and were cultured in DMEM supplemented with 10% fetal calf serum (FCS) and 100 IU/mL of penicillin/streptomycin. Parental and stably-expressing Gag/Pol HEK293 lines were cultured in DMEM supplemented with 10% fetal bovine serum (FBS) and 100 IU/mL of penicillin/streptomycin. Pancreatic ductal epithelial cells (PDECs), derived from female C57BL/6n mice, were cultured as previously described^37,38^: Advanced DMEM/F12 supplemented with 10% FBS (Gibco), 100 IU/mL of penicillin/streptomycin (Gibco), 100 mM Glutamax (Gibco), ITS Supplement (Sigma), 0.1 mg/mL soy trypsin-inhibitor (Gibco), Bovine Pituitary Extract (Gibco), 5 nM T3 (Sigma), 100 μg/mL Cholera toxin (Sigma), 4 μg/mL Dexamethasone (Sigma), 10 ng/mL human EGF (Preprotech). PDECs were cultured on collagen-coated plates (100 μg/mL PureCol 5005, Advanced Biomatrix). Tumor-derived cell lines were generated by an initial mechanical disaggregation/mincing, and tumor fragments were transferred to a solution of type V collagenase (Sigma C9263, 1 mg/mL in HBSS 1X) and incubated at 37 C for 45 minutes. Cell suspensions were supplemented with an equal volume of DMEM 10% FBS and filtered through a 100 μm mesh (BD). Filtered suspensions were centrifuged for 5 min at 1250 rpm, pellets were resuspended in DMEM 10% FBS with penicillin/streptomycin 100 μI/mL and cultured on collagen coated plates (100 μg/mL PureCol 5005, Advanced Biomatrix). Cells were passaged twice to remove non-tumor cells and downstream applications were done with these tumor-derived cell lines.

### Engineering Large Genomic Deletions with MACHETE

To engineer genomic deletions, we developed Molecular Alteration of Chromosomes with Engineered Tandem Elements (MACHETE). The premise behind MACHETE is to give cells that bear the deletion of interest a selective advantage over unedited cells, which is achieved by using a bicistronic cassette consisting of an inducible suicide element and an antibiotic resistance component. This cassette is integrated into the region of interest by CRISPR-Cas9 mediated homology directed repair (HDR). Once cells with stable integration of the cassette are positively selected, they are treated with CRISPR-Cas9 to generate the deletion of interest and edited cells are enriched via negative selection.

#### Identification and in vitro transcription of sgRNAs

We used GuideScan to select optimal sgRNA sequences^65^. For each locus of interest, we identified an sgRNA to introduce the MACHETE cassette by HDR, and sgRNAs to generate the deletion of interest. For the 4C4 locus, we designed two independent sets of guides for each deletion to control for potential off-target effects. We generated sgRNAs as previously described^66^. Briefly, a primer with a T7 adapter and the sgRNA sequence was used to PCR amplify the tracrRNA from a pX330 plasmid. The PCR product was then purified and transcribed using the RNA MAXX In Vitro Transcription Kit (Agilent) to produce the sgRNA. sgRNAs were then column purified (RNA Clean & Concentrator, Zymo Research), eluted in water and aliquoted for later use with recombinant Cas9 (Sigma). Oligos used for sgRNA production are listed in Extended Data Table 4.

#### Generation of HDR donor

To maximize flexibility, MACHETE uses 40-bp homology arms that are introduced by PCR. The locus-specific HDR donors were generated by PCR amplification of the MACHETE bicistronic cassette using a high-fidelity DNA polymerase (Herculase II, Agilent or Q5, NEB). PCR fragments were column purified (Qiagen) and quantified. Primers for targeting are presented in Extended Data Table 4.

#### CRISPR-Cas9–mediated targeting and generation of large genomic deletions

For all CRISPR editing, we used Cas9 ribonucleotide complexes (Cas9 RNPs) with the intended guides, to reduce cloning and limit Cas9 expression. To incorporate Cas9 RNPs and donor PCR, cells were electroporated with the Neon System (Invitrogen) following the manufacturer’s instructions.

##### HDR knock-in of MACHETE cassette

Briefly, cells were trypsinized, washed in PBS once, and counted. Cells were then resuspended in Neon Buffer R and aliquoted for the different electroporation reactions. Each condition used 100 x 10^3^ cells in 10 μL of Buffer R. In parallel, 1 μg of Cas9 (ThermoFisher) and 1 μg of sgRNA were complexed for 15 min at room temperature. For the HDR step, 0.5 μg of donor DNA was added to the Cas9 RNP complex, which was then mixed with the cell aliquot. The cell/RNP/donor mixture was electroporated (1400 V pulse voltage, 20 ms pulse width, 2 pulses). For the selection of cassette knock-in lines, Puromycin (2 μg/mL) was added to the media 48 hours after electroporation. In the case of fluorescence reporters, cells were sorted 48 hours post electroporation (Sony MA900), and further enriched for stable expression one week after this initial sort. Selected cells were expanded to establish the parental KI lines. To validate this initial step, cells were then treated with diphtheria toxin (50 ng/mL) or ganciclovir (10 μg/mL) to assess their sensitivity. On-target integrations were assessed by PCR of gDNA and Sanger sequencing of the product for confirmation. Genotyping primers are provided in Extended Data Table 4.

##### Generation of genomic deletions

KI cells were trypsinized, washed in PBS once, and counted. Cells were then resuspended in Neon Buffer R and aliquoted for the different electroporation reactions. Each condition used 100 x 10^3^ cells in 10 μL of Buffer R. In parallel, 2 μg of Cas9, 1 μg of 5’ flanking sgRNA, and 1 μg of 3’ flanking sgRNA were complexed for 15 min at room temperature. The cell/RNP mixture was electroporated (1400 V pulse voltage, 20 ms pulse width, 2 pulses) and cells were seeded in the absence of selection. 48 hours after seeding, cells were treated with diphtheria toxin (50 ng/mL) or ganciclovir (10 μg/mL) and media was changed every 2 days with ongoing selection. Surviving cells were then passaged and analyzed for the presence of the intended deletion breakpoint, loss of selection cassette, and sensitivity to selection was re-evaluated. Genotyping primers are provided in Extended Data Table 4.

##### Breakpoint high-throughput sequencing

Breakpoint PCRs were purified and sent for amplicon sequencing (Amplicon-EZ, Genewiz) following service guidelines. Raw fastq reads were aligned to the mouse genome (mm10) using bowtie2 with parameters “--local -D 50 -R 3 -N 0 -L 19 -i S,1.0,0.7 --no-unal -k 5 --score-min C,20”.

Aligned SAM reads were processed using custom Rscript to parse the breakpoint location, junction position, direction of the reads, and alignment types. Alignments for a proper break read-pairs have to both aligned to the same breakpoint chromosome; coming from 1 primary and 1 secondary alignment; and breakpoints must be located on opposite sides of the breakpoint junction.

### Flow Cytometry

To assess expression of EGFP, tumor cell suspensions were generated by initial mechanical disaggregation/mincing. Tumor fragments were then transferred to a solution of type V collagenase (Sigma C9263, 1 mg/mL in 1X HBSS) supplemented with soy trypsin inhibitor (Gibco, 0.1 mg/mL) and DNAse I (Sigma, 0.1 mg/mL). Tumor pieces in this disaggregation buffer were transferred to a GentleMACS tube and loaded into the OctoDissociator (Miltenyi). Samples were treated with the mTDK1 program, after which 5 mL of FACS Buffer (PBS 1X, 2% FBS) was added to the sample and the mix was filtered through a 100 μm mesh (BD). The resulting cell suspension was centrifuged and resuspended in FACS buffer. Cells were then treated with Fc block (BD, 1:200 dilution) incubated at 4C for 15 minutes and stained with anti-CD45 AF700 (BD, 1:400 dilution) for 30 min at 4C. Cells were washed and resuspended in FACS buffer supplemented with DAPI (Sigma, 1 μg/mL final). Stained cell suspensions were then analyzed in a MA900 sorter (Sony). EGFP+ cells were analyzed within the CD45-, DAPI-population.

For multi-parametric flow cytometry analysis, tumor cell suspensions were generated as above, and cells were stained with LIVE/DEAD fixable viability dye (Invitrogen) for 30 min at 4C. After this, cells were washed, incubated with Fc block (BD, 1:200) for 15 min at 4 C, and then stained with conjugated antibody cocktails (see Extended Data Table 5 for antibody panels) for 30 min at 4C. After staining cells were washed and fixed (BD Cytofix) for 20 min at 4C, washed again, and stored for analysis. Samples were analyzed in a BD LSRFortessa with 5 lasers, where gates were set by use of fluorescence-minus-one (FMO) controls.

### Animals and In Vivo Procedures

#### Animals

All mouse experiments were approved by the Memorial Sloan-Kettering Cancer Center (MSKCC) Institutional Animal Care and Use Committee (IACUC). Mice were maintained under pathogen-free conditions, and food and water were provided ad libitum. C57Bl/6n and NOD/SCID Il2rg^-/-^ (NSG) mice were purchased from Envigo. Foxn1^nu^ (Swiss nude) mice were purchased from Jackson Laboratory. All mice used were 6 to 8 week-old females.

#### PDAC GEMM-ESC models of Cdkn2a/b loss

Embryonic stem cells (ESCs) bearing alleles to study PDAC were used as previously described^67-69^. Briefly, Ptf1a^Cre/+^; Rosa^26Lox-Stop-Lox rtTA3-IRES-mKate2/+^; Col1a1^Homing cassette/+^ cells were targeted with shRNAs against *Smad4* or *Renilla* luciferase (non-targeting control). Mice were then generated by blastocyst injection of shSmad4 or shRen ESCs, and shRNAs were induced by treatment of the resulting mice with doxycycline in drinking water starting at 5-6 weeks of age. Pancreatic tumor initiation and progression were monitored by palpation and ultrasound imaging, mice were euthanized upon reaching humane endpoints of tumor burden, and samples were collected from primary tumors and metastases (when present). Tumor-derived cell lines were then analyzed by sparse whole genome sequencing and classified according to the type of *Cdkn2a/b* alteration.

#### Orthotopic transplants

For orthotopic transplants of PDEC cells, mice were anesthetized and a survival surgery was performed to expose the pancreas, where either 300,000 cells (for primary MACHETE-edited lines) or 100,000 cells (tumor-derived lines) were injected in the pancreas of each mouse. Mice were then monitored for tumor engraftment (bioluminescence imaging, IVIS) and progression, and were euthanized when overt disease was present in accordance with IACUC guidelines.

#### Experimental metastasis assays

For liver colonization of PDEC cells, mice were anesthetized, and a survival surgery was performed to expose the spleen, where 100,000 cells (tumor-derived lines) were injected in the spleen of each mouse, where the site of injection was then removed and the remainder of the spleen was cauterized (hemi-splenectomy). Mice were then monitored for tumor engraftment and progression and were euthanized when overt disease was present in accordance with IACUC guidelines.

#### Antibody treatments

For IFNAR1 blockade experiments, mice were treated twice per week with either 200 ug i.p. of control IgG (MOPC21 clone, BioXCell) or 200 ug i.p. of anti-IFNAR1 antibody (MAR15A3, BioXCell). For depletion experiments: mice were treated with anti-CD8a antibody (Clone 2.43, BioXCell) or anti-CD4 (Clone GK1.5, BioXCell) with an initial dose of 400 ug i.p., followed by maintenance injections of 200 ug/mouse. Control, IFNAR1 blocking and CD8/CD4 depletion antibody treatments were done twice per week, starting one week prior to the orthotopic transplantation of cells. Treatments were maintained for the entire duration of the experiment. B cell depletion was done by a monthly intravenous injection of anti-CD20 (Clone SA271G2, BioLegend), starting one week prior to orthotopic transplantation of cells.

#### In vivo bioluminescence imaging

Mice were anesthetized and hair over the imaging site was removed. Mice were injected with 200 uL of luciferin i.p. (200 mg/L, PerkinElmer #122799) and bioluminescence was acquired 10 minutes after the luciferin injection in an IVIS Spectrum. For organ imaging, mice were injected with luciferin, euthanized 10 min after the injection, and organ bioluminescence was acquired in an IVIS Spectrum instrument.

#### Imaging and assessment of metastatic burden

Mice meeting endpoint criteria were euthanized and inspected for overt macro-metastatic burden in the abdominal cavity (peritoneum, diaphragm, mesenteric lymph nodes, ovary/fallopian tubes, kidneys, and liver), as well as in the thoracic cavity (lungs and rib cage). Primary tumors and organs were dissected and imaged under a dissection microscope (Nikon SMZ1500) for brightfield and EGFP fluorescence.

### RNA Extraction and cDNA Preparation

RNA was extracted by using the Trizol Reagent (ThermoFisher) following the manufactureŕs instructions. The only modification was the addition of glycogen (40 ng/mL, Roche) to the aqueous phase to visualize the RNA pellet after precipitation. RNA was quantified using a Nanodrop. cDNA was prepared with the AffinityScript QPCR cDNA Synthesis Kit (Agilent) following the manufacturer’s instructions.

### DNA Extraction

Genomic DNA was extracted from cells or tissues using the DNeasy Blood and Tissue Kit (Qiagen) following the manufacturer’s instructions.

### qPCR

For quantitative PCR the PerfeCTa SYBR Green FastMix (QuantaBio), the Taqman Fast Advanced Master Mix (Applied Biosystems), and the Taqman Genotyping Master Mix (Applied Biosystems) were used following manufacturer’s instructions. For qPCR primers and Taqman assays, see Extended Data Table 6.

### Histology

Tissues were formalin fixed, dehydrated and paraffin embedded for sectioning. Hematoxylin / Eosin staining was performed with standard protocols.

### RNA Sequencing, Differential Gene Expression, and Gene Set Enrichment Analysis

Bulk tumor pieces were flash frozen on dry ice and stored at -80C. Tissues were then mechanically disrupted in Trizol and RNA was extracted following manufacturer’s instructions. RNA integrity was analyzed with an Agilent 2100 Bioanalyzer. Samples that passed QC were then used for library preparation and sequencing. Samples were barcoded and run on a HiSeq (Ilumina) in 76 bp SE run, with an average of 50 million reads per sample. RNA-Seq data was then trimmed by removing adapter sequences and reads were aligned to the mouse genome (GRCm38.91; mm10), and transcript counts were used to generate an expression matrix. Differential gene expression was analyzed by DESeq2 ^70^ for 3-5 independent tumors per condition. Principal Components Analysis (PCA) and differentially expressed gene analysis was performed in R, with significance determined by >2 fold change with an adjusted p value < 0.05. GSEA ^71,72^ was performed using the GSEAPreranked tool for conducting GSEA of data derived from RNA-seq experiments (v.2.07) against specific signatures: Hallmark Pathways, Reactome Pathways, and Immune Subpopulations.

### Sparse Whole Genome Sequencing

Low-pass whole genome sequencing was performed on gDNA freshly isolated from cultured cells as previously described ^73^. Briefly, 1 μg of gDNA was sonicated on an E220 sonicator (settings: 17Q, 75s Covaris), and library preparation was done by standard procedure (end repair, addition of polyA, and adapter ligation). Libraries were then purified (AMPure XP magnetic beads, Beckman Coulter), PCR enriched, and sequenced (Illumina HiSeq). Reads were mapped to the mouse genome, duplicates removed, and an average of 2.5 million reads were used for CNA determination with the Varbin algorithm ^74^.

### Human PDAC Transcriptional Analysis

Samples from the COMPASS trial^44,45^ were classified as primary or metastatic disease and further subdivided by status of the 9p21.3 locus: 9pS deletion affecting *CDKN2A/B*, or 9pL deletions affecting *CDKN2A/B* and at least one IFN gene from the linked cluster. 9pS and 9pL samples were then analyzed for differentially expressed genes using DESeq2 and assessed by GSEA for Reactome Pathways^75^, and Immune Subpopulations^42^. As an independent validation of the differences between primary and metastatic PDAC, a previously published dataset^76^ was used to derive differentially expressed genes using DESeq2. Genes downregulated in PDAC metastasis were then analyzed using the Enrichr algorithm^77^.

### scRNA Sequencing

The single-cell RNA-Seq of FACS-sorted cell suspensions was performed on Chromium instrument (10X genomics) following the user guide manual for 3′ v3.1. In brief, FACS-sorted cells were washed once with PBS containing 1% bovine serum albumin (BSA) and resuspended in PBS containing 1% BSA to a final concentration of 700–1,300 cells per μl. The viability of cells was above 80%, as confirmed with 0.2% (w/v) Trypan Blue staining (Countess II). Cells were captured in droplets. Following reverse transcription and cell barcoding in droplets, emulsions were broken and cDNA purified using Dynabeads MyOne SILANE followed by PCR amplification per manual instruction. Between 15,000 to 25,000 cells were targeted for each sample. Samples were multiplexed together on one lane of 10X Chromium following cell hashing protocol ^78^. Final libraries were sequenced on Illumina NovaSeq S4 platform (R1 – 28 cycles, i7 – 8 cycles, R2 – 90 cycles). The cell-gene count matrix was constructed using the Sequence Quality Control (SEQC) package ^79^.

#### Data Pre-processing

FASTQ files were generated from 3 different samples (ΔL, ΔS, α−IFNAR1 ΔS) with three mice pooled together per condition. These files were then processed using the SEQC pipeline^79^ using the default parameters for a 10X single-cell 3’ library. This pipeline begins with aligning the reads against the provided mouse mm10 reference genome and resolving multi-mapping incidents. SEQC then corrects for UMIs and cell barcodes and filters cells with high mitochondrial fraction (>20%), low library complexity (few unique genes expressed), and empty droplets. The resulting count matrix (cell x gene) was generated for each condition as the raw expression matrices.

As each mouse was barcoded with a unique hashtag oligo for each sample, in order to demultiplex the cells, an in-house method known as SHARP (https://github.com/hisplan/sharp) was employed. Labels are assigned to either identify a cell as belonging to a specific mouse or as a doublet/low-quality droplet. The labeled cell barcodes and gene expression matrix were then concatenated together into one count matrix. Most of the downstream analysis and processing was done using the Scanpy software^80^.

#### Data cleanup

We began by filtering for lowly expressed genes defined as those expressed in less than 32 cells in the combined dataset. The resulting count matrix was then normalized by library size (defined as the total RNA counts per cell), scaled by median library size, and log2-transformed with a pseudocount of 0.1 for the combined dataset. For downstream analysis, we first performed dimensionality reduction using Principal Component Analysis (PCA) to obtain top 30 principal components (PCs), chosen based on the decay of associated eigenvalues, computed on the top 4,000 highly variable genes (HVGs). We then computed a k-nearest neighbor graph representation of the cells on the obtained principal components *(n_neighbors = 30)*. We visualized the cells on a 2-dimensional projection using UMAP^81^ based on the implementation in Scanpy (using *min_dist = 0.1* parameter). All the cells from different samples were observed to group together based on their cell type, which indicated that no batch effect was present in the data (**Figure 3A**). The cells were then clustered using PhenoGraph^82^ on the PCA space with *k=30*. We ensured that the clusters were robust to variations around the chosen parameter of *k.* We measured consistency using adjusted rand index (as implemented in the Sklearn package in Python) and observed high degree of consistency for values of *k* around 30. Upon close inspection of the obtained clusters, we observed one cluster that had low CD45 (PTPRC-) and high KRT8+ expression and two other clusters that had low CD45 and high expression of Mitochondrial genes. As such, we decided to remove these clusters from further analysis.

#### IFN response signature

We first sought to broadly understand, on a per cell type basis, the response to IFN activity. We reasoned that to answer this, we ought to identify the genes that are most differential between α−IFNAR1 and control ΔS. As such, we constructed an IFN signature by identifying top 100 differentially upregulated genes in ΔS compared to α−IFNAR1. The differential genes were identified using MAST^83^ and the top 100 genes were averaged on a per cell basis and plotted on the UMAP (**Figure 3C**). Once the signature was constructed, we removed cells from the α−IFNAR1 condition from further analysis in order to directly contrast 1′S and 1′L.

#### Analysis on 1′S and 1′L samples

The count matrix of CD45+ cells from the 1′S and 1′L samples included 15334 cells and 15329 genes, 7774 cells belonging to 1′S and 7560 to ΔL. To ensure that the observed heterogeneity was not impacted by these cell clusters, we re-processed the data using the same parameters as described above. Broad cell types were assigned to these clusters according to the average expression of known markers.

#### CD8+ T cells

We isolated cells identified as CD8+ T cells in order to analyze them separately. For this, the 6,080 cells were sub-clustered using PhenoGraph on top of the first 30 PCs (*k=30*) using 1,500 highly variable genes. Using known markers, these PhenoGraph clusters were then annotated into further subtypes of CD8+ T cells based on the average expression of the markers in each sub-cluster.

#### Milo analysis on CD8+ T cells

We employed Milo^43^ to statistically quantify the changes in abundance of 1′S and 1′L specific cells among the CD8+ T cells subtypes. Milo utilizes nearest-neighbor graphs to construct local neighborhoods (possibly overlapping) of cells and calculates and visualizes differential abundance of cells from different conditions in the obtained neighborhoods. For this analysis, we first constructed a k-nearest neighbor graph (*k=30*) on the first 30 PCs using the *buildGraph* function in Milo. Neighborhoods were calculated using the *makeNhoods* function (*prop=0.1, refined=TRUE*). We used default parameters for *countCells, testNhoods,* and *calcNhoodDistance* in order to calculate statistical significance and spatial FDR correction, and *plotNhoodGraphDA* (alpha=0.5) to visualize the results. The color scale of the logFC uses blue to represent higher abundance of 1′L cells and red to represent higher abundance of 1′S specific cells, and the size of the circle is proportional the number of cells belonging to the neighborhood. We further assigned each neighborhood a cell-type identity if more than 80% of the cells in a neighborhood belonged to a specific CD8+ T subtype, otherwise they are categorized as Mixed.

#### Dendritic cells

Cells annotated as dendritic cells were isolated for further analysis. The 1,134 cells were clustered using PhenoGraph on top 30 principal components (k=30) using 1,500 HVGs. The dendritic cells were further cell typed according to markers from previous studies^84^. The proportion of cells that belong to 1′L and 1′S in each cluster was calculated and plotted.

#### Macrophages

Cells labeled as macrophages (1,788 cells) were isolated. The cells were clustered using PhenoGraph on top 30 principal components (k=30) using 1,500 HVGs. These clusters were analyzed and annotated according to macrophage subtypes based on the differentially expressed genes computed in each cluster compared to the rest of the data using MAST. The proportion of cells that belong to 1′L and 1′S in each cluster was calculated and plotted.

#### B cells

1,204 cells annotated as B cells were selected for. The cells were clustered using PhenoGraph on top 30 principal components (k=30) using 1,500 HVGs. We obtained differentially expressed genes in each B cell sub-cluster using MAST and utilized the results to distinguish distinct populations. The proportion of cells that belong to 1′L and 1′S in each cluster was calculated and plotted.

### General Statistical Analysis

Graphs and statistical analyses for Figures 2, 4, Extended Data Figures 2, 3, 4, 5, 7, and 8 were done with GraphPad Prism. For all experiments n represents the number of independent biological replicates. For Figures 2C, Extended Data Figure 2E, Extended Data 4D, and Extended Data Figure 5A-I differences were evaluated with a two-tailed t-test. For Figure 4A-B, 4H-I, 4J-M, Extended Data 3G, Extended Data 3K, Extended Data 7D, Extended Data 7F, Extended Data Figure 8I-J, differences were assessed by a one-way ANOVA followed by Tukey or Sidak’s multiple comparison test. To assess differences in tumor initiation or metastasis incidence, contingency tables followed by a chi-square test were done for figures: 2A, 2E, 2F, 2H, 2I, 2L, 3K, Extended Data Figure 3F, Extended Data 3M. For survival curves, log rank-test was used to assess significant differences. Differences were considered significant for p values < 0.05, where asterisks represent the level of significance for the analysis used: *, p < 0.05; ** p < 0.01; ***, p < 0.001; n.s. not significant, p > 0.05.

## ACKNOWLEDGEMENTS

We thank Anahi Tehuitzil, Kasia Rybczyk, Sha Tian, and Wei Luan for technical assistance; Francisco J. Sánchez-Rivera, Riccardo Mezzadra, John P. Morris IV, and the rest of the Lowe laboratory for advice and helpful discussions; Camilla Salvagno, Juan Cubillos-Ruiz, Edward R. Kastenhuber, and Charles J. Sherr for advice and discussions; John Erby Wilkinson for pathology analysis. We acknowledge the TCGA datasets generated by the TCGA Research Network; the MSKCC Research Animal Resource Center, Mouse Genetics Core, Small Animal Imaging Core, and Integrated Genomics Operation Core, funded by the NCI Cancer Center Support Grant (CCSG, P30 CA08748), Cycle for Survival, and the Marie-Josée and Henry R. Kravis Center for Molecular Oncology. F.M.B. was supported by a GMTEC Postdoctoral Fellowship, an MSKCC’s Translational Research Oncology Training Fellowship (5T32CA160001-08), and a Young Investigator Award by the Edward P. Evans Foundation. K.M.T. was supported by the Jane Coffin Childs Memorial Fund for Medical Research and the Shulamit Katzman Endowed GMTEC Postdoctoral Fellowship. T.B. is supported by the William C. and Joyce C. O’Neil Charitable Trust, Memorial Sloan Kettering Single Cell Sequencing Initiative. D.A.C. is recipient of the La Caixa Postdoctoral Junior Leader Fellowship (LCF/BQ/PI20/11760006). This work was also supported by MSKCC’s David Rubenstein Center for Pancreatic Research Pilot Project (to S.W.L); the Agilent Thought Leader Program (to S.W.L.); and NIH grant P01CA13106 (to S.W.L.). S.W.L. is an investigator in the Howard Hughes Medical Institute and the Geoffrey Beene Chair for Cancer Biology.

## AUTHOR CONTRIBUTIONS

F.M.B. and K.M.T. conceived the study, designed and performed experiments, analyzed data and wrote the manuscript. Y.-J.H. and A.Z. analyzed WES and RNA-seq data. N.S and R.S. analyzed scRNA Seq data. T.B. analyzed sWGS data. A.N.W., I.D, B.M, G.L., A.P.C., D.A.C and J.S. assisted in experiments. R.C. performed scRNA Seq. D.B-S., C.A.I-D, and F.N. provided critical reagents and data. D.P. provided supervision and critical input on scRNA Seq analysis. S.W.L. conceived and supervised the study and wrote the manuscript. All authors read the manuscript.

## EXTENDED DATA FIGURE LEGENDS

**Extended Data Figure 1.**
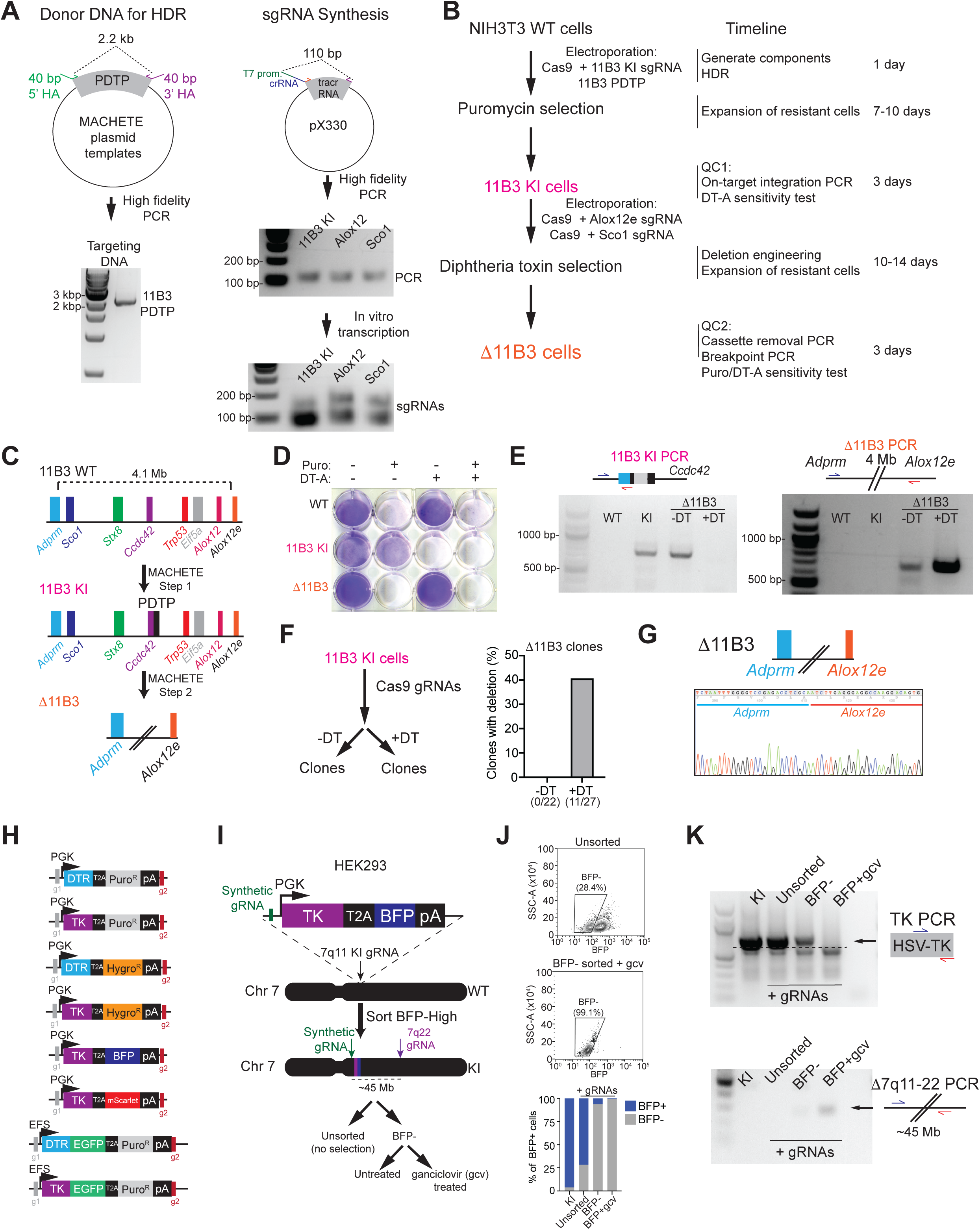
(A) Preparation of donor DNA and sgRNA used for MACHETE-mediated targeting of the 11B3 locus in NIH3T3 cells. (B) Experimental outline and timing for MACHETE-based 11B3 deletion engineering in NIH3T3 cells. (C) Schematic of MACHETE-mediated engineering of a 4.1 Mb deletion at the 11B3 locus. (D) Crystal violet stain of WT, 11B3 KI and 1′11B3 NIH3T3 cells after selection with puromycin (Puro, 2 μg/mL) and/or diphtheria toxin (DT-A, 50 ng/mL). (E) PCR genotyping for the 11B3 KI and 1′11B3 alleles in the indicated NIH3T3 cell lines. (F) (Left) Experimental outline for testing the impact of DT-mediated negative selection on the efficiency of 1′11B3 deletion engineering in NIH3T3 cells. (Right) Clonal analysis of NIH3T3 cells engineered without (-DT) and with (+DT) diphtheria toxin selection. (G) Sanger sequencing of the 11B3 deletion breakpoint confirming the expected deletion. (H) Suite of dual selection cassettes generated for the MACHETE approach. (I) Schematic of MACHETE-mediated engineering of a 45 Mb deletion at the 7q11-22 locus in HEK293 cells. (J) Flow cytometry plots and quantification of BFP+ and BFP-HEK293 cells under the indicated conditions. (K) PCR genotyping for the 7q11 KI and Δ7q11-22 alleles in HEK293 cells under the indicated conditions.

**Extended Data Figure 2.**
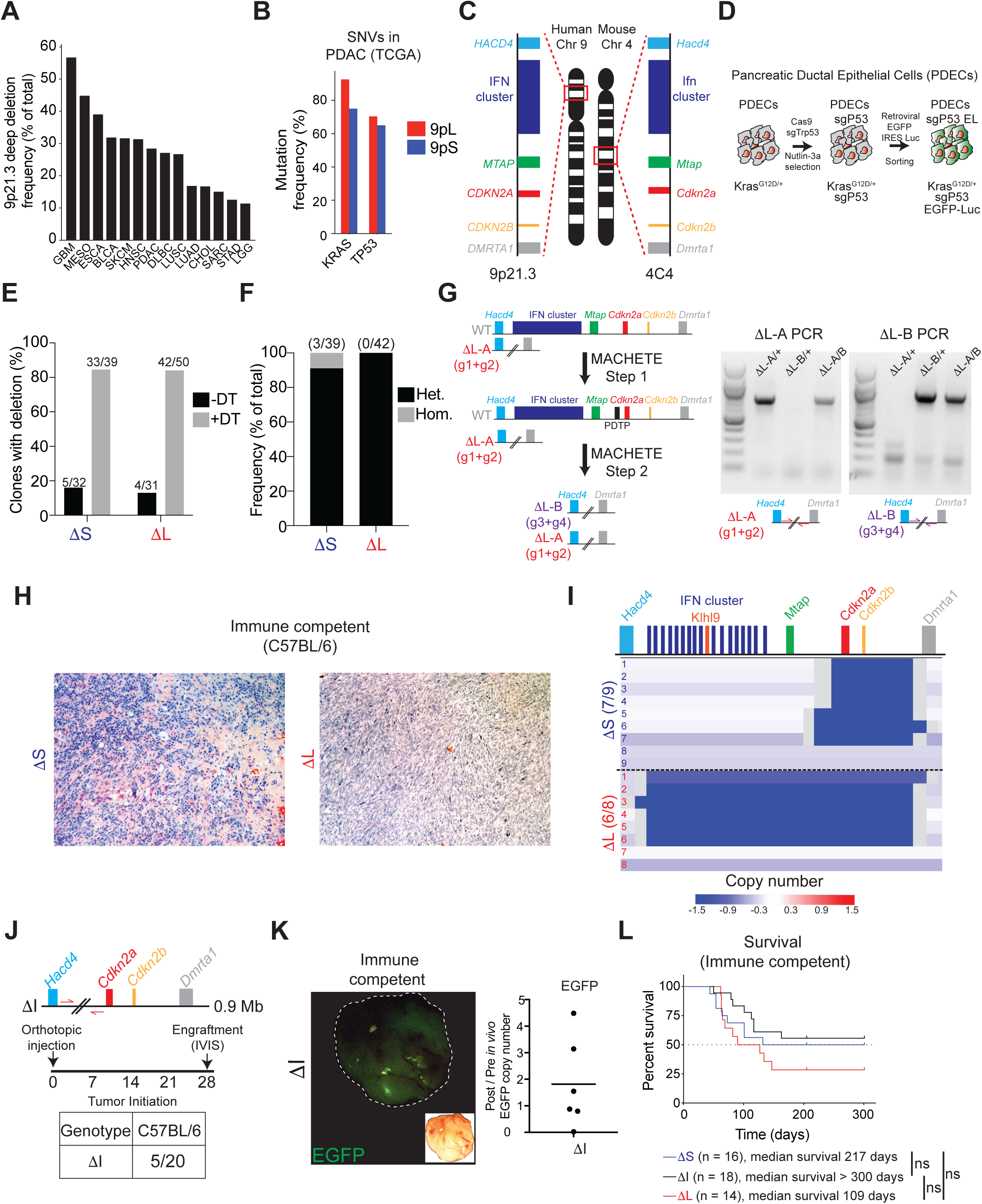
(A) Frequency of deep deletions at the 9p21.3 locus across different types of cancer in the TCGA dataset. (B) Mutation frequency of *KRAS* and *TP53* in 9pL and 9pS PDAC patients in the TCGA dataset. (C) Schematic of the synteny between the human 9p21.3 and mouse 4C4 locus. (D) Schematic of the generation of PDEC sgP53 EL cells. CRISPR-mediated knockout of *Trp53* was done by electroporation of a pX330-sgP53 plasmid followed by treatment with Nutlin-3 (10 μM) to select for *Trp53*-deficient cells. PDEC sgP53 cells were then infected with a retroviral EGFP-Luciferase construct and cells were selected by sorting for EGFP+ expression. (E) Clonal analysis of 1′S and 1′L cells engineered without (-DT) and with (+DT) diphtheria toxin selection. (F) Frequency of heterozygous and homozygous 1′S or 1′L deletions in PDEC cells following MACHETE engineering. (G) (Left) Schematic of iterative editing of cells bearing a heterozygous ΔL deletion, using a distinct set of guides to discern between the different deletions. (Right) PCR genotyping of the distinct 1′L deletion breakpoints. (H) Histology of 1′S and 1′L tumors in C57BL/6 mice. Representative H/E images are shown. (I) sWGS analysis of 4C4 deletion status in 1′S and 1′L tumor-derived cell lines (from C57BL/6 hosts). Deep blue color depicts deletion defined as log2 relative abundance < -2. (J) (Top) Schematic representation of the MACHETE-engineered ΔI allele that removes a 0.9 Mb region downstream of *Hacd4* and upstream of *Cdkn2a*. (Bottom) Engraftment of ΔI cells in C57BL/6 mice one month after injection and measured by bioluminescence. (K) (Left) Representative macroscopic image of a ΔI tumor showing retained EGFP expression at endpoint. Inset shows matched brighfield image. (Right) qPCR analysis for EGFP copy number in the gDNA of tumor-derived (Post *in vivo*) ΔI cell lines from C57BL/6 hosts relative to their parental (Pre *in vivo*) counterparts. Each dot represents an independent cell line. (L) Survival curve of C57BL/6 mice transplanted with 1′S, 1′I, or 1′L tumor cells. Depicted are the number of mice transplanted and the median survival, which showed no statistically significant differences (logrank test).

**Extended Data Figure 3.**
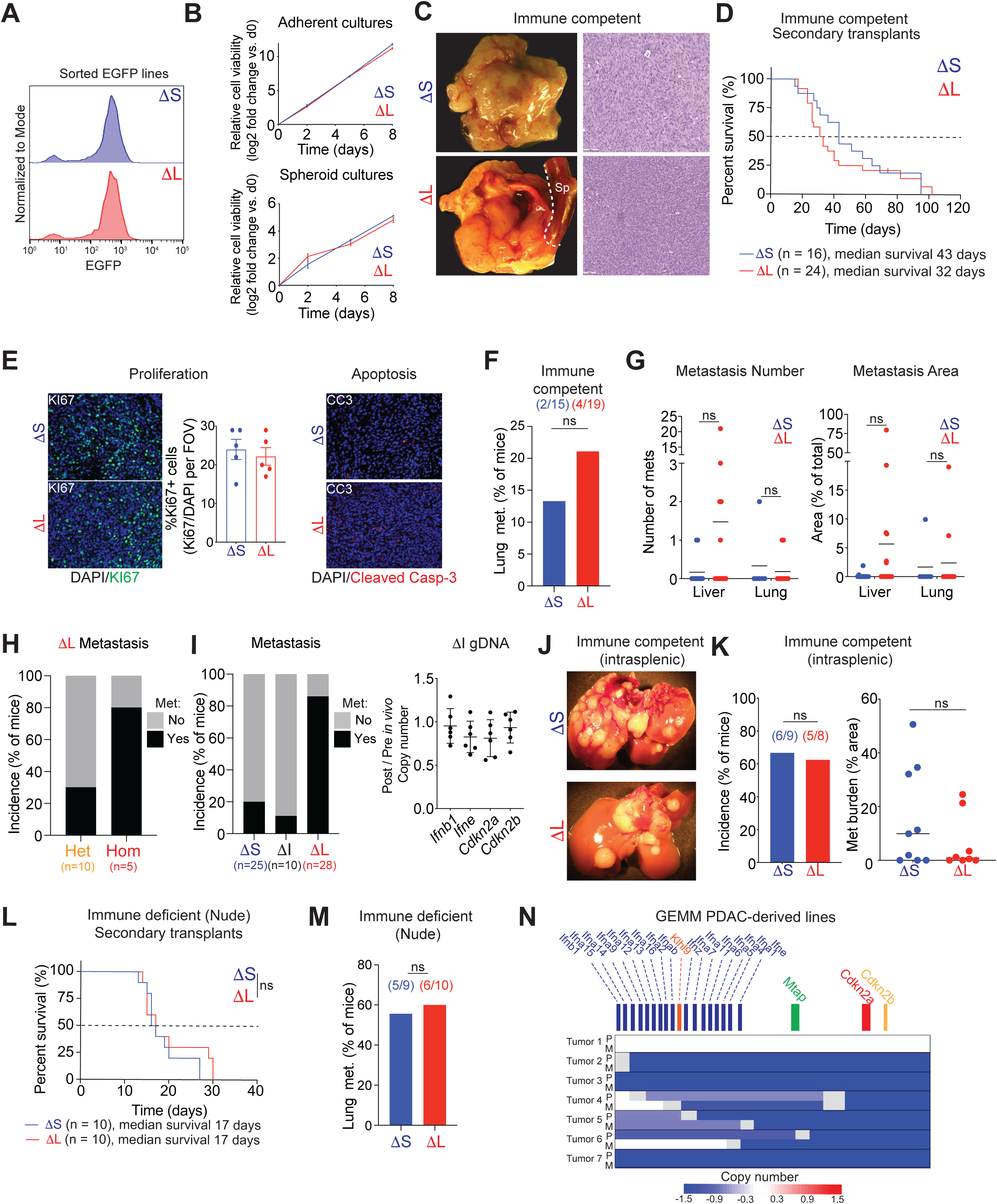
(A) EGFP levels of representative re-sorted tumor-derived 1′S and 1′L cell lines. (B) Growth curves in adherent (top) or suspension (bottom) conditions for 1′S and 1′L cell lines. (C) Macroscopic images (left) and hematoxylin/eosin stain (right) of orthotopic tumors in C57BL/6 mice transplanted with tumor-derived 1′S and 1′L cells. (D) Survival curve of C57BL/6 mice transplanted with tumor-derived 1′S and 1′L cells. (E) Representative images (left) and quantification (middle) of the fraction of Ki67+ cells in 1′S and 1′L tumors. (Right) Representative images of cleaved caspase-3 in in 1′S and 1′L tumors, which showed little to no detectable signal. (F) Lung metastasis incidence in C57BL/6 mice with either 1′S or 1′L tumors. Bars represent fraction of metastasis-bearing mice (specific numbers of independently analyzed mice are noted in parentheses). ns = non-significant, chi-square test. (G) Quantification of the number (left) and relative area (right) of liver and lung metastases in C57BL/6 mice with either 1′S or 1′L tumors. (H) Metastasis incidence in C57BL/6 mice with either heterozygous or homozygous ΔL tumors. (I) (Left) Metastasis incidence in C57BL/6 mice with 1′S, 1′I, or 1′L tumors. (Right) Copy number of *Ifnb1*, *Ifne*, *Cdkn2a*, and *Cdkn2b* in tumor-derived ΔI lines (Post) relative to pre-injection parental 1′I cells (Pre). Each dot represents an independent tumor-derived line. (J) Macroscopic images of liver metastases in C57BL/6 mice after intrasplenic injection of either 1′S or 1′L cells. (K) Relative area of liver metastases in C57BL/6 mice after intrasplenic injection of either 1′S or 1′L cells. (L) Survival curve of Nude mice transplanted with tumor-derived 1′S and 1′L cells. (M) Lung metastasis incidence in Nude mice with either 1′S or 1′L tumors. (N) Analysis of 4C4 deletion status in PDAC GEMM cell lines derived from matched primary tumors (‘P’) and metastases (‘M’). sWGS was used to assess the status of the 4C4 locus. Deep blue color depicts deletion defined as log2 relative abundance < -2.

**Extended Data Figure 4.**
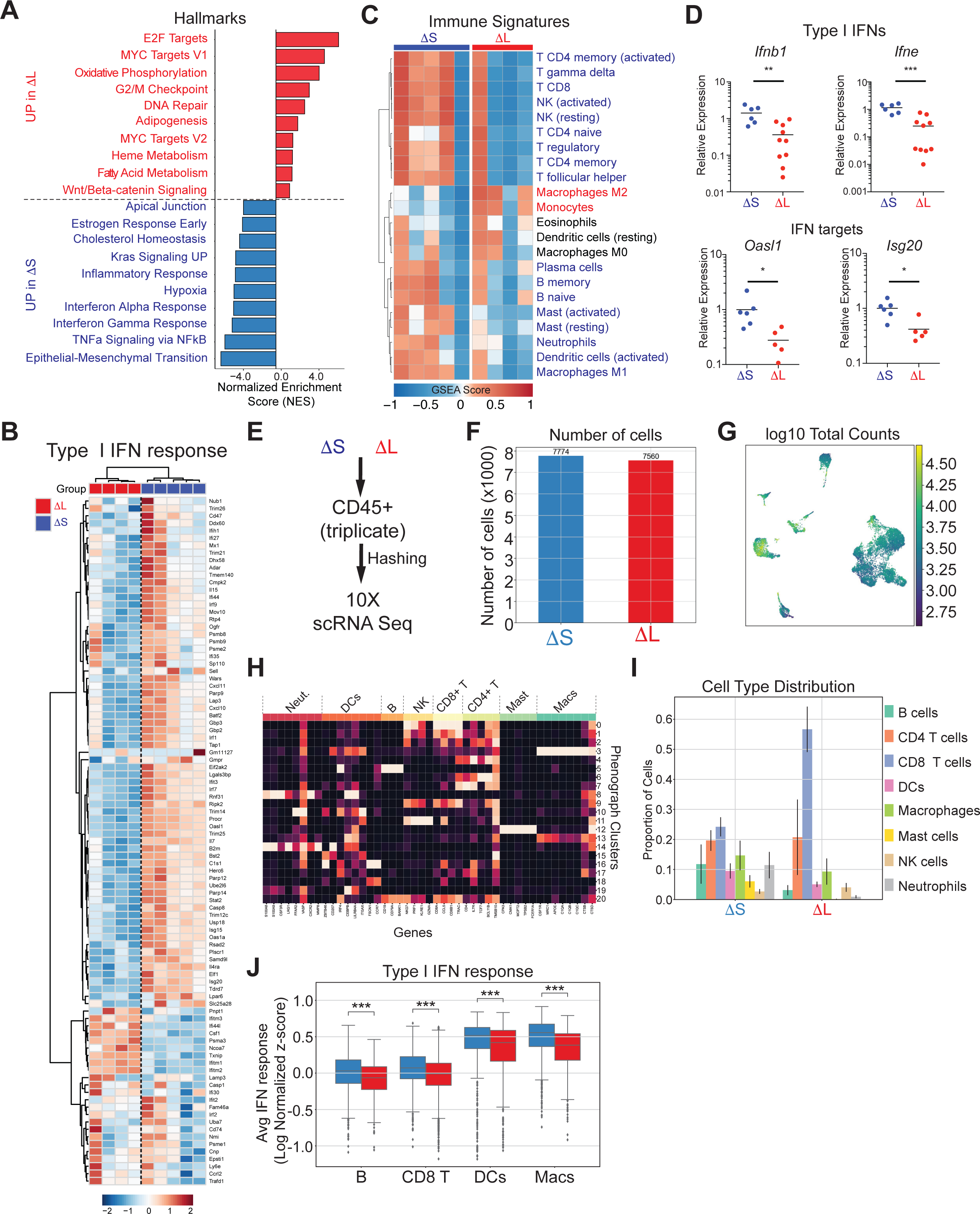
(A) Histogram of GSEA Normalized Enrichment Score (NES) highlighting the top 10 differentially expressed Hallmark gene datasets in 1′S and 1′L tumors. (B) Heatmap of type I IFN response gene expression in 1′S and 1′L tumors. (C) Heatmap of gene expression signatures for distinct immune subpopulations in 1′S and 1′L tumors. (D) Relative mRNA expression of representative type I IFN genes (*Ifnb1*, *Ifne*) or type I IFN targets (*Oasl1*, *Isg20*), measured by RT-qPCR. Each dot represents an independent biological replicate. *p < 0.05, **p < 0.01, ***p < 0.001, two-tailed t-test. (E) Experimental design for scRNA Seq analysis of CD45+ cells. CD45+ cells were sorted from three independent 1′S and 1′L tumors, uniquely labeled by antibody-coupled barcoding, pooled and processed for scRNA Seq analysis. (F) Number of high-quality CD45+ cells recovered from 1′S and 1′L tumors. (G) UMAP of library size per cell. (H) Heatmap of genes used to identify specific subpopulations within CD45+ cells. (I) Distribution of CD45+ cells across different subpopulations in 1′S and 1′L tumors. (J) Average expression of the type I IFN response signature across antigen-presenting populations (B cells, dendritic cells, and macrophages) and CD8+ T cells. ***, p < 0.001.

**Extended Data Figure 5.**
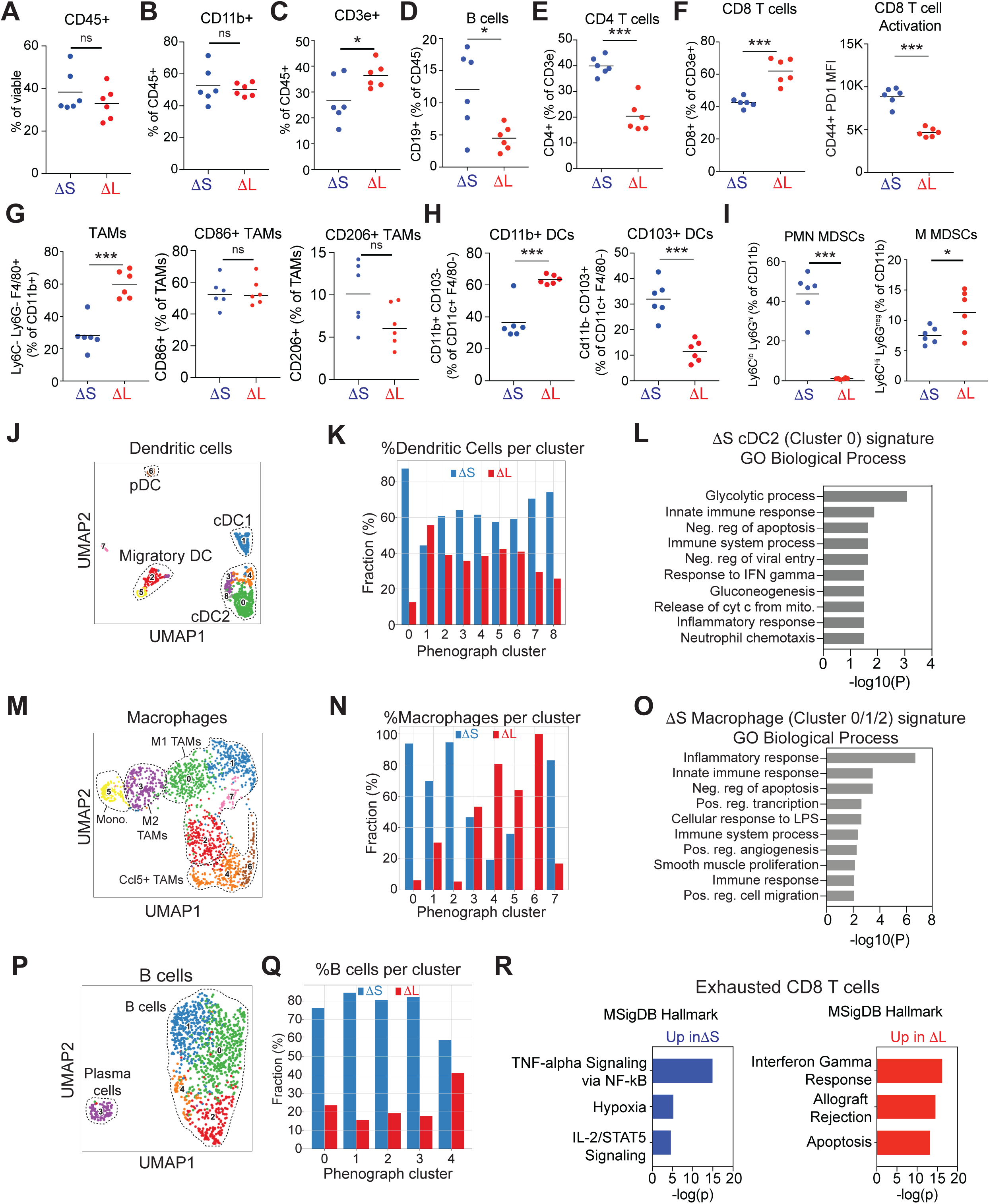
(A-I) Immunophenotyping of infiltrating populations in 1′S and 1′L tumors. Frequency of CD45^+^ cells (A), CD11b^+^ cells (B), CD3e^+^ cells (C), CD19^+^ B cells (D), CD4^+^ T cells (E), CD8^+^ T cells and corresponding PD1 mean fluorescence intensity of CD44^+^CD8^+^ T cells (F), tumor-associated macrophages (TAMs) including CD86+ and CD206+ subtypes (G), CD11b^+^ and CD103^+^ dendritic cell subsets (H), and myeloid-derived suppressor cells (MDSCs) including polymorphonuclear (PMN-MDSCs) and mononuclear (M-MDSCs) subtypes (I). *p < 0.05, **p < 0.01, ***p < 0.001, ns = non-significant; two-tailed t-test. Each dot represents an independent biological replicate. (J) UMAP of dendritic cell phenographs from 1′S or 1′L tumors. Known populations/states are circled. (K) Frequency of dendritic cells across phenographs in 1′S or 1′L tumors. (L) DAVID analysis of Gene Ontology Biological Processes enriched in ΔS-specific dendritic cells. (M) UMAP of macrophage phenographs from 1′S or 1′L tumors. Known populations/states are circled. (N) Frequency of macrophages across phenographs in 1′S or 1′L tumors. (O) DAVID analysis of Gene Ontology Biological Processes enriched in ΔS-specific macrophages. (P) UMAP of B cell phenographs from 1′S or 1′L tumors. Known populations/states are circled. (Q) Frequency of B cells across phenographs in 1′S or 1′L tumors. (R) Enrichr analysis of the top Hallmark Pathways enriched in exhausted CD8+ T cells from 1′S and 1′L tumors.

**Extended Data Figure 6.**
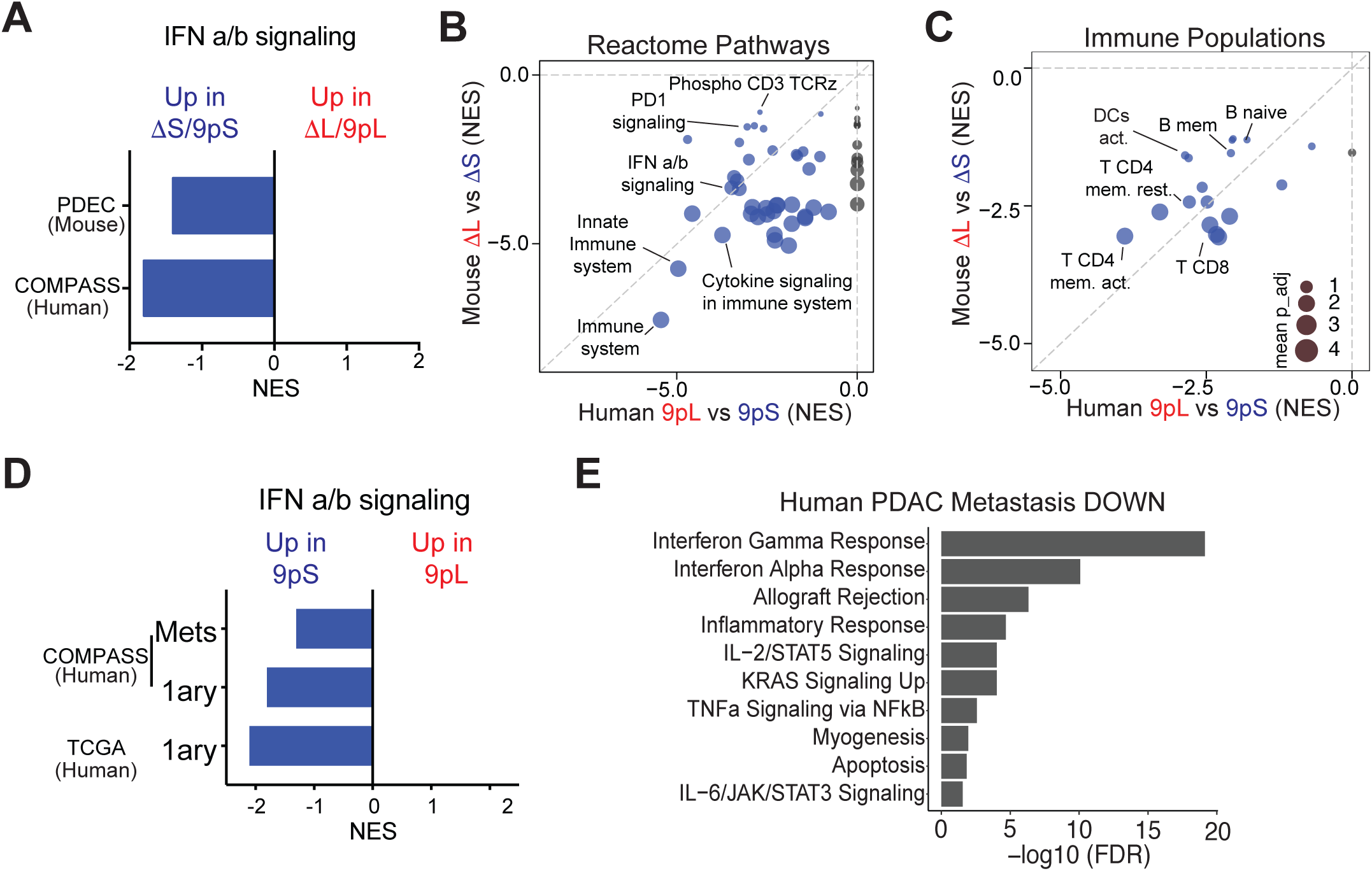
(A) GSEA enrichment scores (NES) of type I IFN signaling in mouse ΔS and human 9pS tumors compared to ΔL and 9pL tumors, respectively. (B) Comparison of GSEA NES scores for Reactome Pathways enriched in mouse ΔS (y axis) and human 9pS tumors (x axis). Highlighted are key pathways and immune populations enriched in IFN-proficient tumors. Circle size represents the adjusted p value. (C) Comparison of GSEA NES scores and Immune populations enriched in mouse ΔS (y axis) and human 9pS tumors (x axis). Highlighted are key immune populations enriched in IFN-proficient tumors. Circle size represents the adjusted p value. (D) GSEA enrichment scores (NES) of type I IFN signaling in human primary or metastatic 9pS tumors compared to 9pL tumors from the COMPASS and TCGA datasets. (E) Hallmark pathways downregulated in human PDAC liver metastases vs. primary tumors. Data from Moffitt et al., 2015^75^.

**Extended Data Figure 7.**
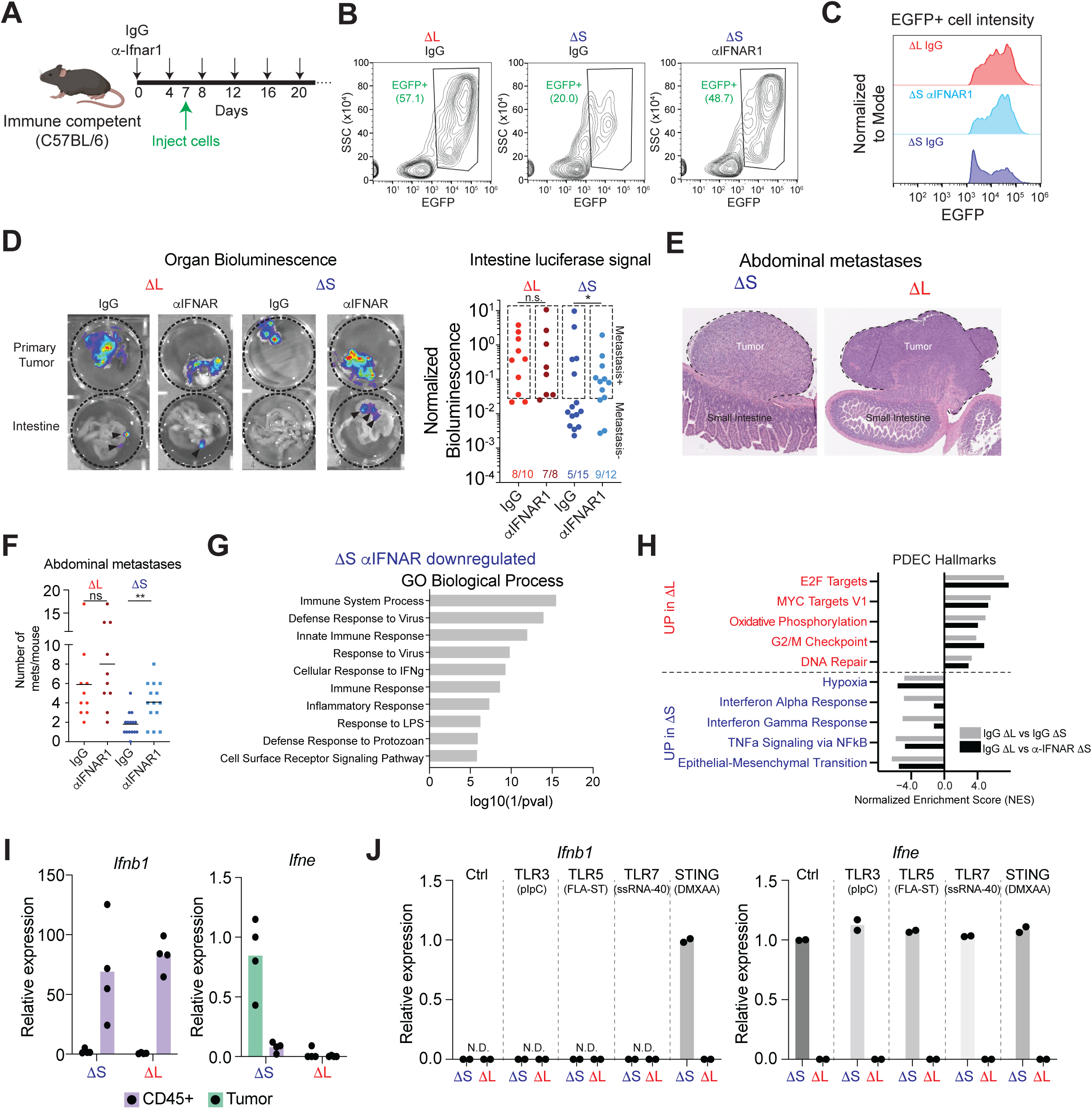
(A) Experimental outline to test the role of type I IFNAR signaling in transplantation experiments. (B) Representative flow cytometry plots of EGFP fluorescence in 1′S or 1′L tumors from C57BL/6 mice treated with IgG or αIFNAR1 antibodies. (C) Representative FACS plots of EGFP+ populations from IgG ΔL, IgG ΔS, or αIFNAR1 1′S tumors. (D) (Left) Representative bioluminescent images of primary tumors and intestines from mice with indicated genotypes of transplanted cells and antibody treatments. (Right) Quantification of all replicates. Boxes indicate the signal threshold for metastasis detection. *p < 0.05, chi-square test. (E-F) Representative H/E images (E) and quantification (F) of mesenteric lymph node metastases in mice with indicated genotypes of transplanted cells and antibody treatments. *p < 0.05, two-tailed t-test comparing IgG vs IFNAR1 blockade in the corresponding cell lines. (G) DAVID gene ontology analysis of α-IFNAR1 downregulated genes in ΔS tumors. Top 10 significant pathways are shown. (H) IFNAR1 blockade specifically affects IFN signaling. NES scores of top 5 UP and DOWN Hallmark categories in tumors comparing 1′L vs 1′S (grey bars, data from Figure 4C) or 1′L vs α−IFNAR1 1′S (black bars). (I) RT-qPCR measurements of mRNA levels for *Ifnb1* and *Ifne* in tumor cells and infiltrating CD45+ cells from 1′S and 1′L tumors. Dots represent independent tumors. (J) qRT-PCR measurements of mRNA levels for *Ifnb1* and *Ifne* in 1′S and 1′L tumor-derived cells after the indicated treatments. Dots represent independent cell lines.

**Extended Data Figure 8.**
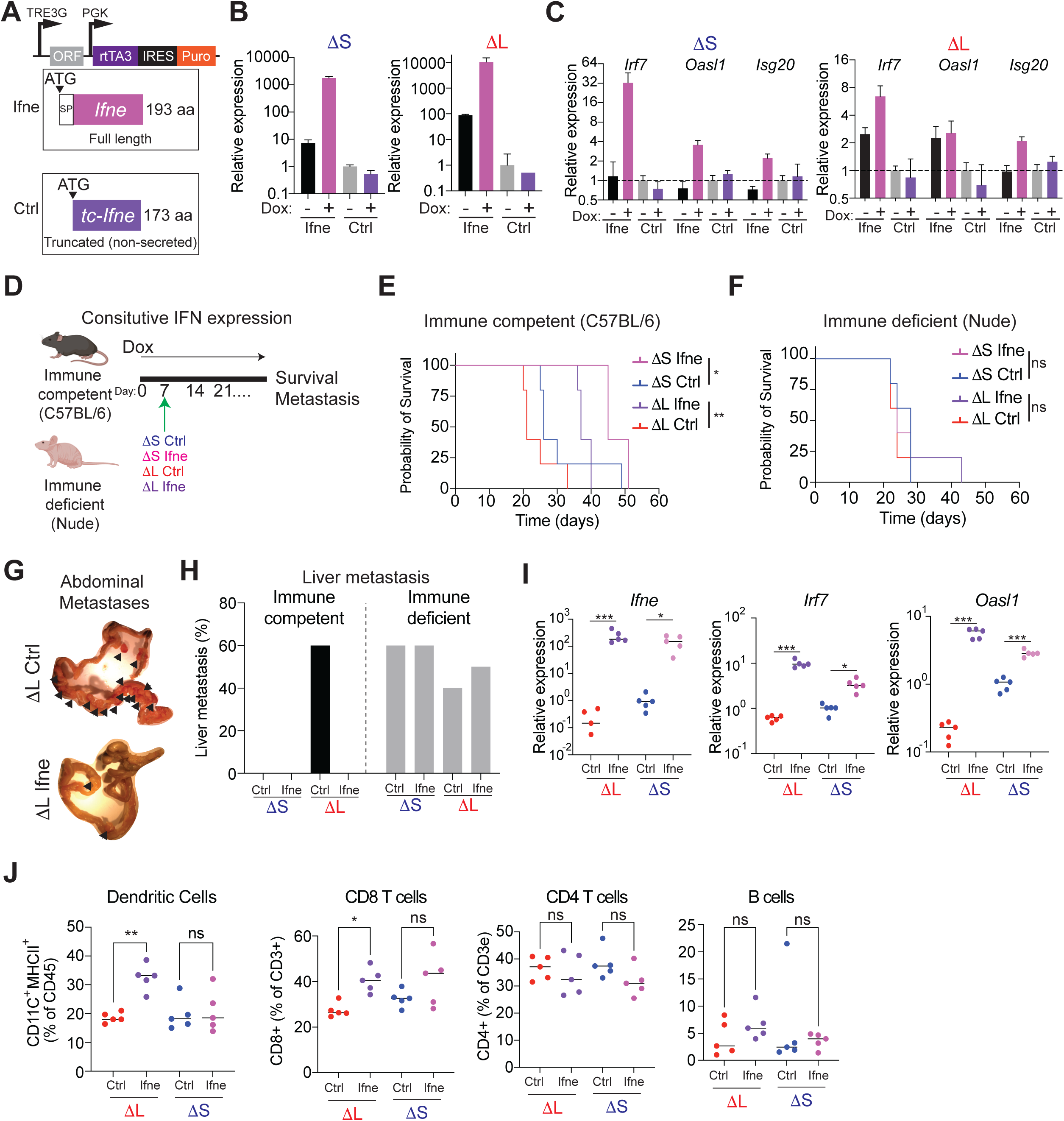
(A) Design of the vector for doxycycline-inducible expression of full-length mouse *Ifne* or a truncated version lacking the signal peptide as control. (B) RT-qPCR of *Ifne* expression in cells cultured -/+ doxycycline (2 μg/mL) for 72 hours. The assay specifically amplifies full-length *Ifne*. (C) RT-qPCR of IFN target genes (*Irf7*, *Oasl1*, *Isg20*) to in cells cultured -/+ doxycycline (2 μg/mL) for 72 hours. (D) Experimental design to test the role of sustained *Ifne* expression in immune competent and immune deficient mice. (E) Survival curve of immune competent mice orthotopically transplanted with Ctrl or *Ifne* overexpressing 1′S and 1′L cells. n = 5 per condition. *p < 0.05; **p < 0.01, log rank test. (F) Survival curve of immune deficient (nude) mice orthotopically transplanted with Ctrl or *Ifne* overexpressing 1′S and 1′L cells. n = 5 per condition. n.s.= non-significant, log rank test. (G) Representative image of an intestine from a mouse with sustained expression of Ctrl or full-length *Ifne* ΔL cells at endpoint. Arrowheads point to macrometastases in the mesentery and intestine. (H) Incidence of overt liver metastasis in immune proficient and deficient hosts transplanted with 1′S or 1′L cells expressing Ctrl or full length *Ifne* (n=5). (I) RT-qPCR of *Ifne*, *Irf7*, and *Oasl1* in tumors from immune competent mice treated with doxycycline for 1 week before tumor analysis. Each dot represents an independent tumor (n=5). *p < 0.05; ***p < 0.001, one-way ANOVA followed by Sidak’s multiple comparison test. (J) Tumor immune infiltration of immune competent mice treated with doxycycline for 1 week before tumor analysis. Frequency of dendritic cells (far left), CD8 T cells (left), CD4 T cells (right), ad B cells (far right) are shown. Each dot represents an independent tumor (n=5). *p < 0.05; **p < 0.01, n.s. = non-significant, one-way ANOVA followed by Sidak’s multiple comparison test.

